# A Humanized Mouse Model of DRPLA Recapitulates Epilepsy and Demonstrates Antisense Oligonucleotide-Mediated Disease Modification

**DOI:** 10.64898/2026.10.12.758592

**Authors:** Qian Wu, Inga Erickson, Stephanie Davidson, Sophie Shonka, Velvet Smith, Bereket Gidi, Kevin M Knox, Tim Yu, Vikram Khurana, Jeff Carroll, Joanna A. Korecka, Melissa Barker-Haliski

## Abstract

Dentatorubral-pallidoluysian atrophy (DRPLA) is an ultra-rare progressive myoclonus epilepsy caused by a glutamine-coding CAG repeat expansion in atrophin-1 (*ATN1*). Mutant *ATN1* (m*ATN1*) expression results in symptoms that inversely correlate to CAG-expansion size: young-onset cases have the most CAG repeats and developmental delay, high-frequency treatment-resistant seizures, and early mortality. We sought to characterize the seizure susceptibility of a fully humanized *ATN1* mouse expressing human m*ATN1* with 112 CAG repeats to then assess therapeutic antisense oligonucleotide (ASO) intervention on seizure-related outcomes. Atn1^Q112/+^ or wild-type (WT) mice were treated with investigational ASO targeting human, CAG-expanded, *ATN1* or saline (P2 and then again at 4 weeks-old) before acute seizure testing, ataxia profiling, and chronic *in vivo* video-EEG recording for spontaneous seizures up to 8 weeks-old. Saline-treated *Atn1^Q112/+^*mice were more seizure prone with early death. However, ASO administration to *Atn1^Q112/+^*mice normalized seizure threshold, decreased ataxia, and protected against early death. *Atn1^Q112/+^* mice exhibited spontaneous behavioral seizures recorded over 2 weeks, which were absent in ASO-treated *Atn1^Q112/+^* and WT mice, with a specific m*ATN1* expression-driven decrease in delta and theta EEG frequencies. *Atn1^Q112/+^* mice are highly seizure prone, validating this model for DRPLA epilepsy. *ATN1* represents a novel epilepsy-related risk factor for therapeutic investigation.

## Introduction

Dentatorubral-pallidoluysian atrophy (DRPLA) is an autosomal dominant neurodegenerative condition characterized by progressive epilepsy, ataxia, chorea, and dementia caused by the expansion of a glutamine-coding CAG trinucleotide sequence in the *atrophin-1* (*ATN1)* gene^1^. It is prevalent in Asia, especially in Japan, with an estimated prevalence of ∼5 per million^2^. DRPLA may also be underdiagnosed in some populations^3; 4^, with accurate gene sequencing only becoming a standard diagnostic tool in recent years. With two clinical phenotypes – juvenile onset and adult onset - clinical diagnosis can be complicated due to the heterogeneous presentation, age of symptom onset, and symptomatic overlap with other spinocerebellar ataxias (SCAs) and Huntington’s Disease (HD). Juvenile-onset DRPLA (onset younger than 20 years) is characterized by intellectual disability and seizures^5^ correlated with the CAG expansion (CAG^EX^)^6^. As such, DRPLA is clinically classified as a progressive myoclonic epilepsy (PME)^7^. The clinical neuropathology of DRPLA includes neuronal loss and astrogliosis in the dentatorubral and pallidoluysian systems, accompanied by neuronal intranuclear ATN1-positive inclusions^7^. Astrogliosis is also present throughout the brain^7; 8^, and neuronal atrophy is observed in the cortex^9^, frontal lobe, and pontine brain areas, especially in juvenile-onset DRPLA^10^. This cortical pathology may account for the observed PME and intellectual disability phenotypes. Life expectancy varies between 8 and 16 years after symptom onset, with no disease-modifying treatments available. While antiseizure medications (ASMs) are the clinical standard-of-care for symptomatic seizures in DRPLA^11^, seizure control is insufficient, and clinical prognosis is ultimately poor. Preclinical studies of the cellular and molecular mechanisms contributing to *ATN1* CAG^EX^ disease pathology and seizure outcomes could thus improve quality of life for DRPLA patients and their caregivers, as well as potentially inform targeted interventions^7^.

*ATN1* is a transcriptional coregulator involved in nuclear receptor signaling^7^, and may control neuronal precursor cell differentiation during development^12^. *ATN1* knockout mice are viable and do not exhibit obvious neurologic phenotypes, albeit minimal brain histopathological analysis has been thus far conducted^13^. On the other hand, human data imply that *ATN1* could have an essential role in CNS development, with heterozygous loss-of-function *ATN1* variants appearing to be under selective constraint (pLI=1.0 in gnomAD), and reports of a non-progressive neurodevelopmental disorder, CHEDDA syndrome, caused by missense variants in the *ATN1* HX domain^14^. With a clear role in neuronal development, yet ambiguous continued cellular expression into adulthood^7; 8^, the function of *ATN1* in driving seizure susceptibility and onset of spontaneous recurrent seizures (SRS) in the juvenile versus adult brain is relatively unknown.

Hindering the clinical development of impactful anticonvulsant and disease-modifying interventions for DRPLA has been the limited preclinical investigation into seizure-related phenotypes and seizure-associated outcomes in this PME. Expression of CAG^EX^ *Atn1* in mice and *Drosophila* causes neurodegenerative changes consistent with human DRPLA^13^. Limited anecdotal reports in a serendipitously expanded mouse DRPLA model suggested the presence of SRS^15^, albeit no comprehensive study of behavioral or electrographic seizure phenotypes was included. More recently, a novel humanized mouse model of DRPLA has emerged^16^, removing a major obstacle to the discovery and advancement of impactful disease-modifying therapeutic options for DRPLA. This mouse DRPLA model exhibits motor dysfunction including ataxia, circadian activity disruptions, and p62-immunoreactive inclusions in the cortex, thalamus, and olfactory nucleus^16^. Functional dysregulation of calcium and neuropeptide signaling^17–19^ and altered autophagy^20^ further support the link between the expression of toxic CAG^EX^ *Atn1* and disease pathogenesis. However, the comprehensive analysis of PME-related seizure symptom presentation in DRPLA models has not yet been reported, leaving a major gap in modeling DRPLA symptom progression, as well as establishing whether targeted suppression of *ATN1* expression can modify seizure symptoms of DRPLA.

Antisense oligonucleotides (ASOs) are promising therapeutic options for deleterious gain-of-function genetic variants, the presumed mechanism for CAG expansion disorders like DRPLA. ASOs are approved or are in clinical trials for several CNS disorders^21–25^, including FDA breakthrough designation for a rare, drug-resistant pediatric epilepsy, Dravet syndrome^26^, as well as n=1/few investigational new drug (IND) and pre-IND programs^27; 28^, like those targeting ataxia-telangiectasia^29^. We thus hypothesized that m*ATN1* expression in our newly developed humanized DRPLA model^16^ could also reproduce the clinical PME features, including SRS onset, reduced seizure threshold, and seizure-related hippocampal neuroinflammation. Further, we hypothesized that pathological disease markers, seizure susceptibility, and SRS presentation would be attenuated or reversed by ASO infusion. This is the first feasibility study to demonstrate that ASO administration in a humanized DRPLA model may delay or even halt the progression of key disease-related seizure features, providing the rationale for future lead compound differentiation studies to advance new gene therapy treatments for DRPLA. This study thus extends our prior in vivo DRPLA investigation^16^ to establish that this model demonstrates clear behavioral and pathological similarities to clinical PME^16; 30^. This model provides a suitable platform to study the cellular mechanisms underlying DRPLA seizure pathology and to evaluate novel antiseizure and anti-epileptogenic therapies for this and other PMEs.

## Methods

### Mice and investigational ASO administration

*Atn1^Q112/+^* mice were generated via CRISPR/Cas9 editing, as described^16^. These mice have one complete mouse *Atn1* allele replaced with human *ATN1*, which has 112 pure CAG repeats inserted into exon 5. C57BL/6NTac oocytes were used for *in vitro* fertilization with cryopreserved *Atn1^Q112/+^* sperm, which were then implanted into recipient dams that were shipped to the University of Washington (UW). Viable pups born at UW were used in these experiments.

For the minimal clonic threshold studies with prototype ASMs, WT age-matched male and female C57Bl/6N mice were acquired from Charles River. This study was not testing the effects of sex hormones on behavior or protein expression; thus, we did not track estrous cycles of females or the dominant/subordinate status of males. Mice were housed on a 14:10 light cycle (on at 6 am; off at 8 pm) in ventilated cages in corncob bedding in a manner consistent with the Guide for the Care and Use of Laboratory Animals. All animal work was approved by the UW Institutional Animal Care and Use Committee (protocol 4387-01) and conformed to ARRIVE guidelines ^31^. Animals were provided free access to irradiated chow (PicoLab Rodent Diet 20-5053), filtered water, and enrichment, except during periods of behavioral manipulation. All experimental minimal clonic seizure threshold testing was performed between 7 am and 7 pm.

The investigational *ATN1*-targeting gapmer ASO was administered via intracerebroventricular (ICV) injection at postnatal day 1-2 (30 ug) and then again at 4 weeks of age (200 ug) into both the *Atn1^Q112/+^* and WT mice, with sterile phosphate buffered saline (PBS) used as control. Details on the ASO were previously published.^16^ Total group sizes for all treatments and seizure tests are included with each figure.

### Ataxia scoring

Motor deficits in mice were evaluated at 5 and 8 weeks-old using a previously published protocol for spinal cerebellar ataxia phenotype severity^32^. Briefly, mice were assessed using the behavioral tests including cage ledge walking, hindlimb clasping, gait, and kyphosis. Each test consisted of 3 trials scored on a scale of 0 to 3, with 3 being the most severe phenotype. The cohort completed a single trial of all four tests in a fixed, sequential order before repeating the sequence for subsequent trials. Scores across the three trials were averaged for each test and then summed to generate a composite score ranging from 0 to a maximum of 12 for each tested animal. All scores were assigned by a researcher blinded to the animal genotype and treatment condition.

### Minimal clonic seizure threshold test

Minimal clonic seizures were evoked with a 60 Hz 0-14 mA 0.2 sec electrical stimulation delivered to anesthetized corneas of awake, lightly restrained 4 month-old WT mice for ASM response studies (Figure 2A and 2B), or 5–8-week-old WT and *Atn1^Q112/+^* mice for the DRPLA model studies (Figure 2C and 2D). Stimulations were delivered every 3-5 days to WT and *Atn1^Q112/+^* mice until at least two points between the limits of 0% and 100% of mice with seizures were established, with seizure threshold as median convulsant current and 95% confidence intervals determined by a Probit regression, model as previously reported^33; 34^. Seizure outcomes were scored with the binary presentation (Y/N) of vibrissae twitching, ventral neck flexion, and forelimb clonus^35-37^, with a minimum of n=4 mice per electrical current. For ASM response studies in WT mice, the investigational ASM was formulated in 0.5% methylcellulose and delivered at the doses tested at the previously established time of peak pharmacological efficacy by the intraperitoneal route (i.p.)^38^.

### Subcutaneous PTZ (scPTZ) test

Subcutaneous (sc) administration of the convulsant, pentylenetetrazol (PTZ), produces clonic seizures in rodents ^39; 40^. The scPTZ test uses a GABA_A_ receptor antagonist and is a clinically validated model of generalized myoclonic seizures and spike-wave discharges^40^, which has contributed to the identification of approved therapies for absence epilepsy^41^. Susceptibility to myoclonic jerk (twitch) and a clonic seizure within 30 minutes of subthreshold scPTZ administration (30 mg/kg in 0.9% saline) was assessed in male and female WT and *Atn1^Q112/+^* mice (n=5-17/group) at 5-to 8-weeks of age. Mice were placed in isolation cages to minimize stress and observed for 30 minutes for the presence or absence of a clonic twitch and/or seizure. An episode of clonic spasms, approximately 3-5 seconds, of the fore-and/or hindlimbs, jaws, or vibrissae was taken as the endpoint. Animals that did not meet this criterion were considered “protected”. Brains of all mice were then collected 2 hours later and flash-frozen to assess brain region activation in response to seizure activity, consistent with our methods for the immediate early gene, c-Fos^38^.

### EEG implant procedure

A subset of saline and ASO-treated *Atn1^Q112/+^* and saline-treated WT mice were enrolled for prospective continuous video-EEG (vEEG) monitoring of potential SRS presentation beginning at 6 weeks-old and lasting until 8 weeks-old (Supplemental Figure 1). Animals were implanted for vEEG monitoring at 5 weeks of age. On the day of surgery, WT and *Atn1^Q112/+^* mice (5 weeks-old; n=4-6/group) were anesthetized with isoflurane (3% induction; 1.0–2.5% maintenance in oxygen) and secured in a stereotaxic frame. Body temperature was maintained at 37 °C, and ophthalmic ointment was applied to prevent corneal drying. Local lidocaine/bupivacaine (1–2 mg/kg) was administered subcutaneously at the surgical site for local anesthesia. The scalp was shaved and sterilized with alternating betadine and 70% ethanol scrubs. A midline incision was made to expose the skull, and connective tissue was removed to clearly visualize Bregma and Lambda. A prefabricated EEG headmount (Pinnacle, #8201-3) was positioned along the midline using Bregma and Lambda as anatomical reference points and secured with tissue adhesive. Pilot holes were created using a sterile needle (# 27), and four stainless steel screws (two anterior 0.10″, Pinnacle #8209; two posterior 0.12″, Pinnacle #8212) were inserted into the skull. Silver epoxy (Pinnacle #8226) was applied to ensure electrical contact, and the assembly was secured with dental acrylic (Supplemental Figure 2). After surgery, anesthesia was discontinued, and mice recovered on a warming pad until ambulatory. Bacon-flavored carprofen tablet (0.5 mg per mouse; Bio-serv MD150) was provided one day prior to surgery and again on the surgery day for analgesia. Implanted mice were individually housed to recover for 1 week before commencing *in vivo* prospective monitoring for SRS activity beginning at 6 weeks-old.

### EEG recording, review and scoring methods

Continuous vEEG recordings were performed in mice using the Sirenia® acquisition system (Pinnacle Technology, Lawrence, KS) with Pinnacle preamplifiers (#8406). A cross-over study design was performed to maximize EEG data acquisition over a 15-day period, enabling the monitoring of 24 total mice using 12 recording chambers operated in alternating sessions (Supplemental Figure 1). Seizure analysis was conducted offline. Seizure identification followed a semi-automated, unbiased workflow. Candidate seizure events were first flagged using automated rodent seizure detection software (ASSYST, Kaoskey), which identified epileptiform activity based on the frequency characteristics of single spikes (processing window = 200 ms; frequency range = 15–25 Hz)^42^. Seizures were operationally defined as spike–wave discharges with amplitudes at least twofold greater than baseline awake activity and a duration of ≥10 s. All software-detected events were then independently reviewed by a trained investigator blinded to experimental conditions and subsequently confirmed by a second trained investigator, also blinded to experimental conditions. Behavioral seizure severity was graded according to a modified Racine scale: stage 0.5, electrographic seizures without observable behavioral manifestations; stage 1, jaw clonus or chewing movements; stage 2, head bobbing and tremor; stage 3, unilateral forelimb clonus; stage 4, bilateral forelimb clonus and/or rearing; and stage 5, stage 4 behaviors accompanied by loss of posture, impaired balance, or tonic extension. Outcome measures included seizure burden (combination of seizure severity x frequency) and behavioral seizure duration. Group sizes for vEEG monitoring were WT saline (n=4; 2 male, 2 female); *Atn1^Q112/+^*saline (n=6; 3 male, 3 female); *Atn1^Q112/+^* ASO-treated (n=5; 3 male, 2 female).

A *post hoc* power band analysis was performed using EEG recordings processed with ASSYST software. Because of the crossover study design (Supplemental Figure 1), mice in cohort 1 had an approximately 15-day interval between the first and last EEG recording sessions, whereas mice in cohort 2 had an approximately 9-day interval between recordings. To improve comparability between cohorts, we calculated daily power change (%) to normalize the data. The normalization equation is presented in Figure 5I. To assess power across different frequency bands, 24 sampling segments were manually selected to represent the entire 24-hour recording period. The selection criterion consisted of one sampling segment from each whole hour (e.g., 1 pm, 2 pm,…, 1 am), resulting in a total of 24 segments per animal. To minimize the potential influence of seizure activity on the analysis of background brain activity during disease progression, sampling segments containing seizure events were avoided. Specifically, if a seizure occurred at the designated sampling time point, the sampling window was shifted 15 minutes later (e.g., from 1:00 pm to 1:15 pm). Only one mouse required this adjustment due to a single seizure event occurring during a designated sampling window. Each sampling segment consisted of a 10-second EEG recording and was analyzed automatically by ASSYST software across a frequency range of 0.1–100 Hz with 1000 FC (frequency characteristic) samples. The resulting data were exported into spreadsheets, and individual frequency bands were subsequently separated for statistical analysis.

### Cryosectioning and Immunofluorescence for NeuroInflammation and Neuronal Activation

Brain tissue was collected in 4% paraformaldehyde in 1xPBS and refrigerated at 4°C. 24 hours after collection, tissue was moved into 30% sucrose with 0.1% sodium azide until cryosectioning of hemibrains (Leica DM1850). The left hemispheres from scPTZ mice were sagittally sliced for cFos immunofluorescence; the right hemispheres from scPTZ and EEG mice were coronally sliced for neuroinflammatory markers (GFAP, Iba-1). Both hemispheres were sliced at a thickness of 30 µm, mounted on Fisher Superfrost Plus microscope slides (4 sections/slide), and stored at 4 C until processing.

The protein product of the immediate early-gene cFos was detected via immunofluorescence on sagittally sliced tissue to assess seizure-induced brain region activation by scPTZ, consistent with our published methods^38^. Briefly, slides were processed with cFos antibody (1:1000; Abcam, ab222699) under 200 µL coverwells in a 1% bovine serum albumin (BSA), 1% goat serum, and 0.03% Triton X-100 in 0.1 M PBS solution for 1 hour at 37 C oven, then 4 C overnight. The following day, coverwells were removed, slides were washed (3 × 5 min in 0.1 M PBS) before being incubated with goat anti-rabbit IgG H&L 555 secondary antibody (1:1000; Abcam, ab150078) and anti-NeuN conjugated to Alexa Fluor 488 (1:300; Sigma, MAB377X) in a 1% goat serum and 0.03% Triton X-100 in 0.1 M PBS solution light protected for 2 hours at room temperature. The slides were again washed and then coverslipped with Prolong Gold with DAPI (ThermoFisher, P36935).

Coronal and sagittal sectioned tissues were used to assess gliosis stratified by genotype and ASO treatment in mice aged 5-(scPTZ) and 9-weeks-old (end of vEEG). The GFAP protein was used to assess astroglial activation, consistent with our established methods^43-45^. The NeuN antibody was conjugated to Alexa Fluor 488 (1:300; Sigma, MAB377X) and GFAP antibody was conjugated to Cy3 (1:1000; Millipore, C9205). The Iba-1 and CD68 proteins were used as markers of microgliosis and microglial activation, respectively. Iba-1 immunofluorescence followed our published methods^43^. After cryosectioning, slides for visualization were washed (3 × 5 min) in 0.3% Triton X-100 in 0.1 M PBS at room temperature. Slides were then incubated in a 5% BSA blocking solution in 0.3% Triton X-100 in 0.1 M PBS under coverwells in a humid chamber for 1 hour. Anti-Iba-1 (1:1000; Wako, 019-19741) and anti-CD68 (1:1000; Neobiotechnologies, 968-MSM3-P0) primary antibodies were applied under 200 µL coverwells in a 0.3% Triton X-100 in 0.1 M PBS solution overnight at 4 C. The following day, coverwells were removed and slides washed (3 × 5 min) in 0.3% Triton X-100 in 0.1 M PBS before being incubated with goat anti-rabbit secondary conjugated to Alexa Fluor 555 (1:1000; Abcam, ab150078), goat anti-mouse secondary conjugated to Alexa Fluor 647 (1:2000; ThermoFisher, A-21235), and anti-NeuN conjugated to Alexa Fluor 488 (1:300; Sigma, MAB377X) in a 1% BSA and 0.3% Triton X-100 in 0.1 M PBS solution in a humid chamber for 2 hours at room temperature. The slides were again washed 3 × 5 min with 0.3% Triton X-100 in 0.1 M PBS before being coverslipped with Prolong Gold with DAPI.

Slides were imaged on an upright fluorescent microscope (Leica DM6 B) with constant acquisition settings (100 ms exposure, 1x gain). Coronally-sliced hippocampal (CA1, CA3, and dentate gyrus) and sagittally sliced cerebellar regions were imaged on tissue immunoreactive to Iba-1, NeuN, GFAP, and CD68 with a 20x objective. Extent of cFos immunopositive signal was rated in sagittal brain sections according to a qualitative scale representing the density of signal in each region (0 = none, 1 = diffuse [1-10 cells], 2 = light [11-30 cells], 3 = moderate [31-80 cells], and 4 = heavy [81+ cells]) indicating the density of neuronal activity, as previously described^38; 46^. The percent field area of NeuN positive signal was measured to assess neuronal density, as previously described^44^.

Images were scored by two investigators blinded to treatment and genotype. Any image with inconsistent scores between investigators was re-assessed by both researchers, and the agreed upon score was used. Raw greyscale 16-bit images from the Iba-1, NeuN, and GFAP channels were utilized to measure the percent field area of immunolabeling-positive cells using ImageJ Fiji (1.54p). Then Iba-1-positive cells co-labeled with CD68 were identified using the colocalization plugin (https://imagej.net/ij/plugins/colocalization.html), and the area of the co-labeled signal measured using ImageJ.

### Statistics

Post-weaning body weight was compared by one-way ANOVA. Post-weaning survival and the latency to twitch in the scPTZ test were compared to age-matched PBS-infused WT mice by Kaplan-Meier survival plot, with p<0.05 considered significant. Ataxia scores were compared by two-way ANOVA between genotype and treatment conditions. Seizure threshold in the minimal clonic test was determined as median convulsant currents and 95% confidence intervals using the Probit regression model^47^, consistent with our established approach in other genetically modified mouse models^33^. If present and detected by vEEG, the number of SRS was compared within groups by two-way ANOVA test (genotype x ASO). vEEGs were compared within groups during each 72-hour bin (5 pm - 8 am).

Percentage changes in power were analyzed using the Kruskal–Wallis test followed by Dunn’s multiple comparisons test. Immunofluorescent signal (cFos rating score and the percent field area of GFAP, NeuN, Iba-1, and Iba-1 co-labeled with CD68) in tissue from mice in the scPTZ test was compared with a two-way ANOVA (genotype x treatment). Immunofluorescent signal (GFAP, NeuN, Iba-1, and CD68/Iba-1 colabeling) in tissue from mice with vEEG monitoring was compared with one-way ANOVA test. All statistical analyses were conducted with GraphPad Prism v10.0 or later, with p<0.05 considered statistically significant, except for median convulsant current. For seizure threshold, statistical difference was defined as non-overlapping 95% confidence intervals.

## Results

### Atn1^Q112/+^ mice exhibited premature death, lessened body weight gain, and worsened ataxia that is rescued by ASO intervention

As we have earlier reported, the novel, fully humanized mouse DRPLA model exhibits significantly worsened overall outcomes consistent with clinical DRPLA^16^. To further characterize the pathological epilepsy-related features of this novel disease model, we herein extended our earlier study to quantify the effects of m*ATN1* expression on body weight change, overall survival, and ataxia from birth to 9 weeks-old (Figure 1) to determine whether this model mirrors the worsened trajectory of PMEs, like juvenile onset clinical DRPLA^11^. While genotype was not determined until after mice were weaned, ASO infusion at P2 and again 4 weeks of age did not adversely affect overall survival to weaning age of all animals in the study (p>0.05; Figure 1A). Separating the treatment conditions by genotype in animals that reached weaning age (>21 days old) revealed a significant treatment x genotype interaction on overall survival of weaned mice, with saline-treated *Atn1^Q112/+^*mice exhibiting significantly reduced overall survival to 8-weeks old (Figure 1B). Notably, ASO infusion to suppress m*ATN1* expression led to significantly improved overall survival such that ASO-treated *Atn1^Q112/+^*mice showed no change in overall survival relative to saline-and ASO-treated WT mice (Figure 1B), whereas saline-treated *Atn1^Q112/+^* mice had substantially reduced survival versus saline-treated WT mice (X^2^= 7.59, p=0.006). Thus, m*ATN1* lowering via investigational ASO delivery at P2 and 4 weeks of age normalized long-term survival of *Atn1^Q112/+^*mice through 9 weeks of age.

**Figure 1.**
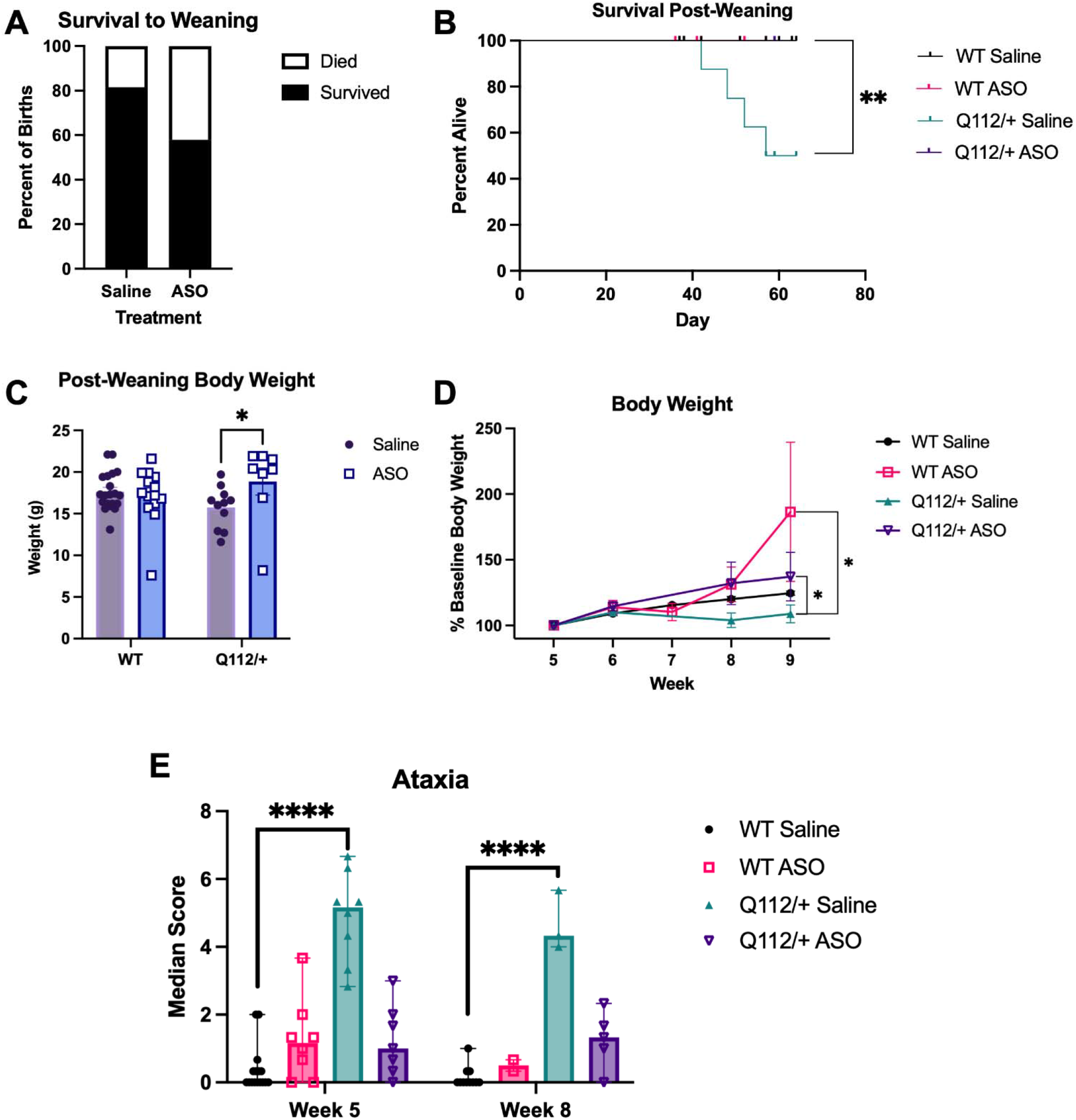
The effect of m*ATN1* expression and suppression by an investigational ASO was evaluated in *Atn1^Q112/+^* mice from birth until 5-9 weeks-old, with euthanasia no later than 9 weeks old. A) ASO infusion at P2, the pre-weaning period, did not adversely influence overall survival relative to saline-treated mice prior to weaning age of 3 weeks (t=2.36, p=0.14; saline n = 38; ASO n = 38). B) Genotype was not determined until weaning age, and all weaned pups were again treated with the investigational ASO at 4 weeks old before commencing behavioral testing. Genotype was determined for each treatment group at weaning. There was significant mortality in saline-treated *Atn1^Q112/+^*mice over the study monitoring period, whereas *Atn1^Q112/+^* mice treated with the investigational ASO had normalized survival post-weaning (WT saline n=20, WT ASO n=14, *Atn1^Q112/+^*saline n=11, *Atn1^Q112/+^* ASO n=8; ** indicates p=0.0024). C) Post-weaning average body weight +/- SEM taken at 5 weeks old showed a significant genotype x treatment interaction (F (1, 49) = 4.076, p=0.049) with post-hoc differences between *Atn1^Q112/+^* mice treated with saline versus the investigational ASO (* indicates p=0.032). D) Age-related increase in body weight gain during the 5–9-week monitoring period showed that saline-treated *Atn1^Q112/+^* mice weighed significantly less than saline-treated WT mice (*p=0.010). However, body weight deficits were rescued by ASO treatment, as *Atn1^Q112/+^* mice treated with the investigational ASO were significantly different from saline-treated *Atn1^Q112/+^* mice (*p=0.049). E) Saline-treated *Atn1^Q112/+^* mice aged 5-9 weeks-old demonstrated worsened motor coordination and ataxia versus saline-treated WT mice. However, this ataxia was rescued by ASO administration. Median with range, * indicates p<0.05.

There was a significant genotype x treatment interaction on post-weaning body weight between experimental groups at 5-weeks-old (F (1, 49) = 4.076, p=0.049), with post-hoc differences between *Atn1^Q112/+^* mice treated with saline versus the investigational ASO (p=0.032; Figure 1C). In addition to ASO treatment effects on long-term survival, body weight changes of *Atn1^Q112/+^* were also tracked from weaning until study conclusion to define whether ASO infusion could improve growth. Indeed, consistent with the overall improved survival over the study monitoring period, *Atn1^Q112/+^* mice generally exhibited reduced body weight gain during the maturation period from 5 to 9 weeks-old, which was rescued by early interventional ASO infusion (Figure 1D). There was no significant difference between ASO-treated *Atn1^Q112/+^* and ASO-treated WT mice in overall body weight change from baseline weights (Figure 1D), but saline-treated *Atn1^Q112/+^* mice had significantly lower overall body weight gain. Notably, ASO-treated WT mice showed a marked increase in overall growth rate relative to saline-treated WT mice (Figure 1D), despite being no different at the 5-week-old monitoring period (Figure 1C). Thus, suppression of m*ATN1* in *Atn1^Q112/+^* mice improves overall body weight gain during the juvenile and young adult development period to match levels in WT mice.

As a putative progressive ataxia model, we also sought to define the progressive worsening of motor coordination deficits and ataxia through a simple behavioral battery assay^32^. Saline-treated *Atn1^Q112/+^*mice exhibited a consistently worsened ataxia score across all testing ages compared to littermate controls (5-and 8-weeks-old; Figure 1E), consistent with our earlier studies^16^. Notably, investigational ASO infusion significantly improved ataxia scores in *Atn1^Q112/+^* mice to match levels observed for WT mice. Thus, early-life ASO infusion led to significantly improved motor coordination and balance in early adulthood, supporting the disease-modifying effect of m*ATN1* suppression.

### Atn1^Q112/+^ mice exhibit altered acute forebrain seizure threshold that is normalized by early mATN1 lowering

The minimal clonic test is a minimally invasive electrically evoked acute seizure model that can establish the integrity and efficiency of neuronal circuits in a rodent with genetic or pharmacological manipulation^33; 34^. We first established the acute anticonvulsant response to the ASMs that are most routinely prescribed in DRPLA cases^11^ to determine how these ASMs would alter minimal clonic test seizure threshold in 4-month-old seizure-naïve male and female WT C57BL/6N mice (Figure 2A and 2B). Notably, only high dose carbamazepine and valproic acid significantly increased the current at which an acute seizure occurred in both male and female WT C57BL/6N mice, whereas the ASMs that have been reported to beneficially control seizures in clinical DRPLA cases^11^ did not acutely influence seizure threshold in WT C57BL/6N mice at the doses tested. Thus, the minimal clonic test is useful to acutely evaluate seizure susceptibility^34^, but mechanistically distinct ASMs did not universally increase the electrical current at which an acute seizure would occur. This differentiated effect of ASMs in the minimal clonic test in WT mice is consistent with findings in other seizure threshold tests, including the thermally evoked seizure threshold test in an *Scn1a^+/-^*mouse model^48^ and the intravenous pentylenetetrazol test ^49^.

**Figure 2.**
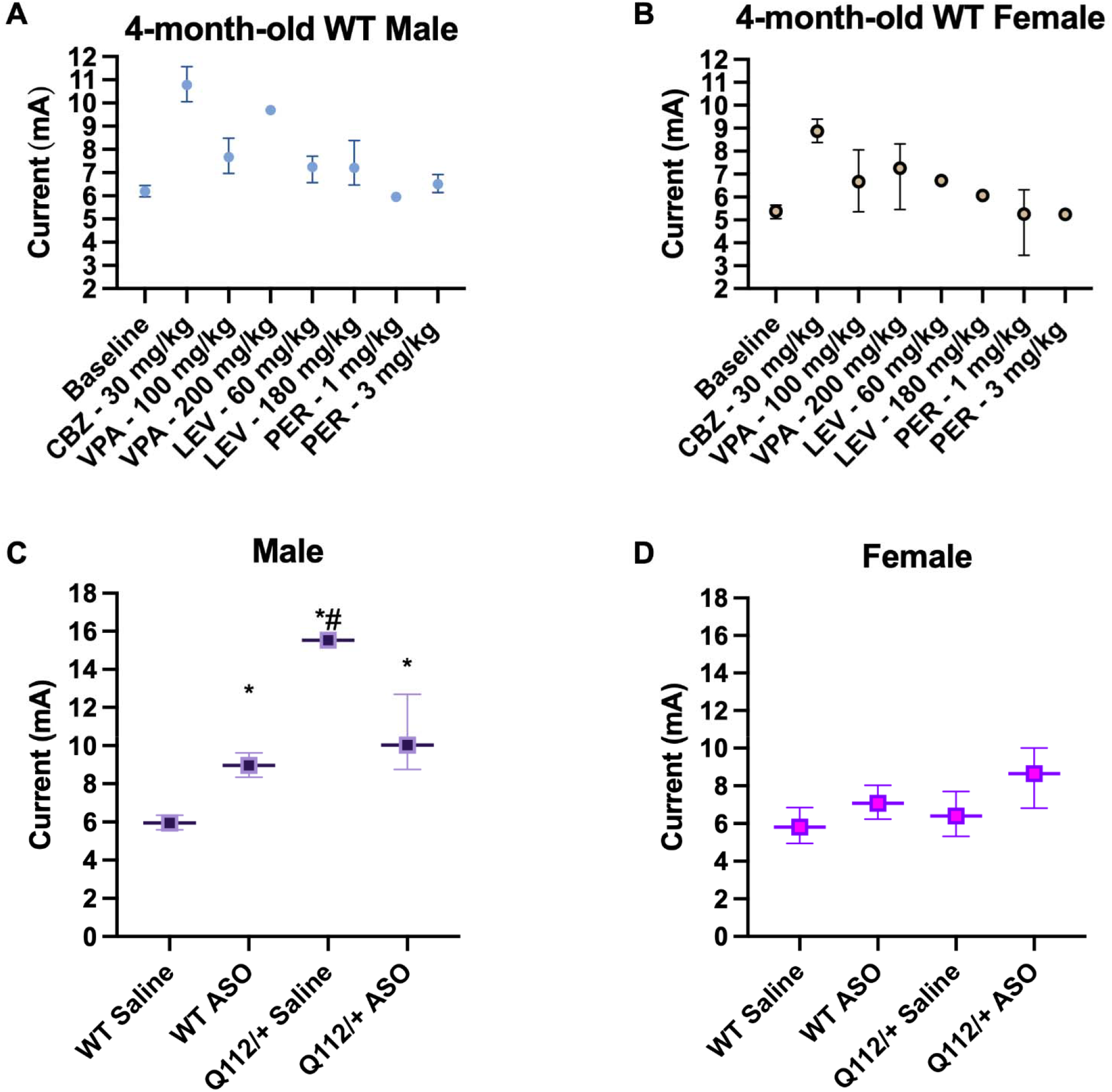
The efficacy of commonly used antiseizure medicines for DRPLA, including carbamazepine (CBZ), valproic acid (VPA), levetiracetam (LEV), and perampanel (PER), was assessed in wild-type (WT) A) male and B) female C57BL/6J mice aged 4-months old relative to define how the commonly prescribed ASMs for DRPLA affect minimal clonic seizure threshold in WT mice. WT mice were housed at UW for a minimum of 1 month prior to behavioral testing using a staircase estimation procedure to define the median convulsive current (CC50) in mA with 95% confidence intervals, as determined by a Probit regression model. Then, the effect of infusion of the investigational ASO or saline was assessed in C) male and D) female WT and *Atn1^Q112/+^* littermates aged 5-8 weeks-old in the minimal clonic test of forebrain seizures. The investigational ASO was delivered at P2 and again at 4-weeks of age before in vivo seizure testing. In all panels, mice were allowed a minimum of 1 hour acclimatization to the testing room prior to behavioral seizure testing. Data are presented as CC50 in mA with 95% confidence intervals, as determined by a Probit regression model. Across all panels, current steps included n=3-10 mice/current, with a minimum of 4 currents tested to establish at least two points between the limits of 0% and 100% of animals with evoked seizures. Wherever 95% confidence intervals do not overlap, differences are statistically significant (* indicates difference from saline-treated WT; # indicates difference from ASO-treated *Atn1^Q112/+^*).

We selected the minimal clonic seizure threshold test to initially establish whether mATN1 expression would in any way alter seizure propensity in young adult mice because it is a rapid and minimally invasive evoked seizure test. In neurologically normal or seizure-naïve animals, the necessary mechanisms for forebrain and brainstem-gated seizure onset function separately, but with genetic manipulation, such as m*ATN1* expression, this neurocircuitry becomes interconnected and focal seizure threshold can be reduced as the neuronal network becomes more efficient^50^. Notably, the minimal clonic test does not include the severe, generalized tonic-clonic seizure of the more common electrically evoked maximal electroshock test^39; 51^ that is routinely used for epilepsy therapy discovery, but which can cause sudden death in WT C57BL/6 mice^52^. As such, the minimal clonic test was selected to assess acute seizure susceptibility and determine the effects of ASO administration on this SRS and epilepsy biomarker in the highly sensitive young adult *Atn1^Q112/+^* mice. We uncovered significant shifts in the post-electrical stimulation behavioral manifestations dependent on the *Atn1^Q112/+^*mutation, which led to significant increases in the perceived seizure threshold exclusively in male *Atn1^Q112/+^* mice (Figure 2C). Female *Atn1^Q112/+^* mice did not demonstrate significant changes in seizure threshold. While it seems counterintuitive that male *Atn1^Q112/+^* mice exhibited elevated seizure threshold, the significant ataxia (Figure 1E) and notable motor deficits^16^ likely influenced the binary blinded seizure scoring of bilateral forelimb clonus and ventral neck flexion in *Atn1^Q112/+^*male mice.

Nonetheless, the investigational ASO delivered to male *Atn1^Q112/+^* mice normalized the seizure threshold vs saline-treated WT mice (as indicated by non-overlapping confidence intervals; Figure 2C), while there was no significant change with ASO infusion in female *Atn1^Q112/+^* mice. There was also significant change in seizure threshold in WT mice similarly treated with the investigational ASO, suggesting that *Atn1* modulation may represent an altogether novel anticonvulsant strategy in mice.

This initial study established that the investigational ASO could normalize acute seizure studies in *Atn1^Q112/+^* mice and compelled further acute seizure susceptibility and chronic SRS monitoring to fully characterize the translational potential of the humanized *Atn1^Q112/+^* mouse model for DRPLA.

### Atn1^Q112/+^ mice are more prone to chemically induced acute myoclonic seizures, an effect that is not rescued by early ASO administration

To further confirm the effect of *Atn1^Q112/+^* variant on myoclonic seizure threshold, we used the well-established scPTZ mouse model of spike-wave discharges that is routinely used for early ASM discovery and is particularly sensitive to ASMs that are effective against generalized nonconvulsive (absence, myoclonic) seizures^40^. Doses of >85 mg/kg PTZ typically evoke seizures in >97% of male WT mice^40; 53^; responses are characterized by first a myoclonic twitch followed by a clonic seizure within 30 min of chemoconvulsant administration. However, for this test with *Atn1^Q112/+^* mice that were already known to be more seizure-prone than WT mice based on the minimal clonic threshold testing, we elected to use a significantly reduced PTZ dose (35 mg/kg, i.e. a dose below the convulsant dose for 97% of normal mice^40; 53^) anticipating significantly greater response likelihood in *Atn1^Q112/+^*mice. In response to low-dose PTZ, few WT mice treated with either saline or ASO exhibited a myoclonic twitch during the 30 minutes after PTZ administration (Figure 3A). Conversely, all saline-treated *Atn1^Q112/+^*mice exhibited rapid onset of myoclonic twitch within 15 minutes of PTZ delivery (Figure 3A), with a significantly reduced twitch latency compared to saline-treated WT mice (X^2^=15.1, p<0.0001). Early ASO administration did not completely rescue threshold to the acute chemically evoked myoclonic twitch in *Atn1^Q112/+^* mice, as ASO-treated *Atn1^Q112/+^* mice were not significantly more prone to myoclonic twitch than saline-treated *Atn1^Q112/+^* mice during in the 30-minute observation period (Figure 3A).

**Figure 3.**
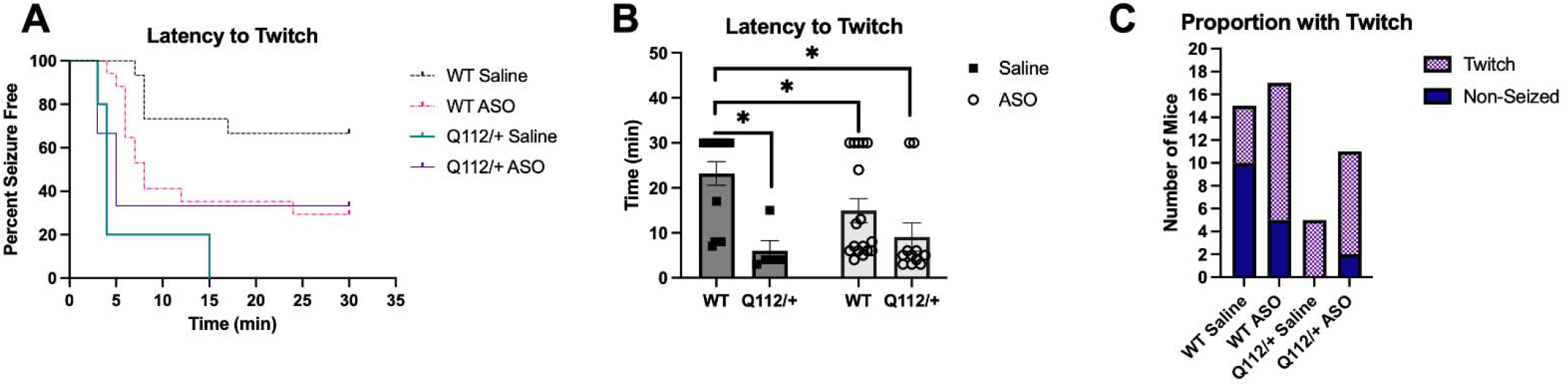
The effect of the investigational ASO or saline was assessed in C) male and D) female WT and *Atn1^Q112/+^* littermates aged 5-to 8-weeks-old in the subcutaneous pentylenetetrazol (scPTZ) model of acute myoclonic and spike-wave discharge seizures. In all panels, mice were allowed a minimum of 1 hour acclimatization to the testing room prior to behavioral seizure testing. A) Infusion of a sub-threshold dose of PTZ (35 mg/kg) does not typically evoke myoclonic twitch or clonic seizures in a majority of WT male and female mice within a 30 min window; however saline-treated *Atn1^Q112/+^* littermates were significantly more prone to the chemoconvulsant effects of this compound. However, ASO infusion did not significantly rescue this effect in *Atn1^Q112/+^* mice. ASO-treated WT mice were more prone to scPTZ infusion. B) The average latency to first myoclonic twitch following a PTZ infusion in saline-treated *Atn1^Q112/+^* mice was significantly reduced (p<0.05), but ASO infusion in WT and Atn1^Q112/+^ mice did not meaningfully rescue twitch latency. C) The proportion of tested mice with myoclonic twitch was markedly affected, with *Atn1^Q112/+^* mice being generally more sensitive to the proconvulsant effects of PTZ (p=0.02). This response was not rescued by the investigational ASO administration. Samples sizes were: WT saline n = 17; WT ASO n = 20; *Atn1^Q112/+^* saline n = 5; *Atn1^Q112/+^* ASO n=11. * indicates p<0.05.

However, when the average latency to first twitch (Figure 3B) and proportion of total mice with a myoclonic twitch (Figure 3C) were compared between treatment groups, *Atn1^Q112/+^* mice were consistently more likely to experience this behavior in response to a low PTZ dose, reflecting an increased sensitivity to the proconvulsant effects of PTZ. All saline-treated *Atn1^Q112/+^* mice (5/5; 100%) exhibited a myoclonic twitch during the 30-minute observation period, whereas only 5/15 (33%) of WT mice exhibited this response (p=0.014; Figure 3C). In the ASO-treated *Atn1^Q112/+^* mice group, 9/11 (81%) of mice had PTZ-induced myoclonic twitch and this response also differed from saline-treated WT mice (p=0.02). Thus, *Atn1^Q112/+^* mice were generally more sensitive to the proconvulsant effects of PTZ and this response was not rescued by the investigational ASO administration. Investigational ASO administration to WT mice also modestly reduced latency to first twitch (p=0.017; Figure 3A) and proportion of mice with twitch relative to saline-treated WT mice (p=0.035; Figure 3C), suggesting a seizure phenotype-specific effect of m*ATN1* suppression. Combined with the minimal clonic response (Figure 2), these scPTZ data indicated that *Atn1^Q112/+^*mice exhibit reduced myoclonic seizure susceptibility long-term, but that early ASO treatment did not acutely increase seizure threshold, compelling the more comprehensive long-term monitoring for potential SRS.

### Atn1^Q112/+^ mice exhibit epileptiform EEG activity and spontaneous seizures

During our initial model characterization and ataxia scoring^16^, we observed suspected spontaneous seizure-like behaviors, which typically occurred in response to handling, justifying the further seizure threshold and susceptibility testing (Figures 2 and 3). To fully characterize the seizure phenotypes associated with this novel humanized DRPLA model, we thus aimed to prospectively and objectively determine whether the *Atn1^Q112/+^* mice had electrographic seizures and confirm SRS via video-EEG monitoring over in a 15-day period (Figure 4). Blinded vEEG review revealed unambiguous spontaneous focal (Figure 4A) and generalized spontaneous seizures (Figure 4B) exclusively in *Atn1^Q112/+^* mice, further quantified by EEG activity between genotypes (Figure 4C-E), and consistent with our prior initial suspicions of seizure-like behavioral events. Notably, only saline-treated *Atn1^Q112/+^* mice demonstrated SRS with clear electrographic features; no saline-treated WT and ASO-treated *Atn1^Q112/+^* mice experienced SRS during the 15-day crossover monitoring period from 6-8 weeks of age (Figure 4D). Representative spectrograms from *Atn1^Q112/+^* and WT mice further confirmed marked differences in EEG power during the monitoring period (Figure 5). Due to the limited study size and prior minimal clonic threshold data to suggest minimal changes from saline-treated WT mice, we did not prospectively monitor ASO-treated WT mice via vEEG. Nonetheless, our chronic monitoring of *Atn1^Q112/+^* mice via vEEG confirmed the suspected epileptic phenotype with both electrographic and motor features, consistent with human DRPLA.

**Figure 4.**
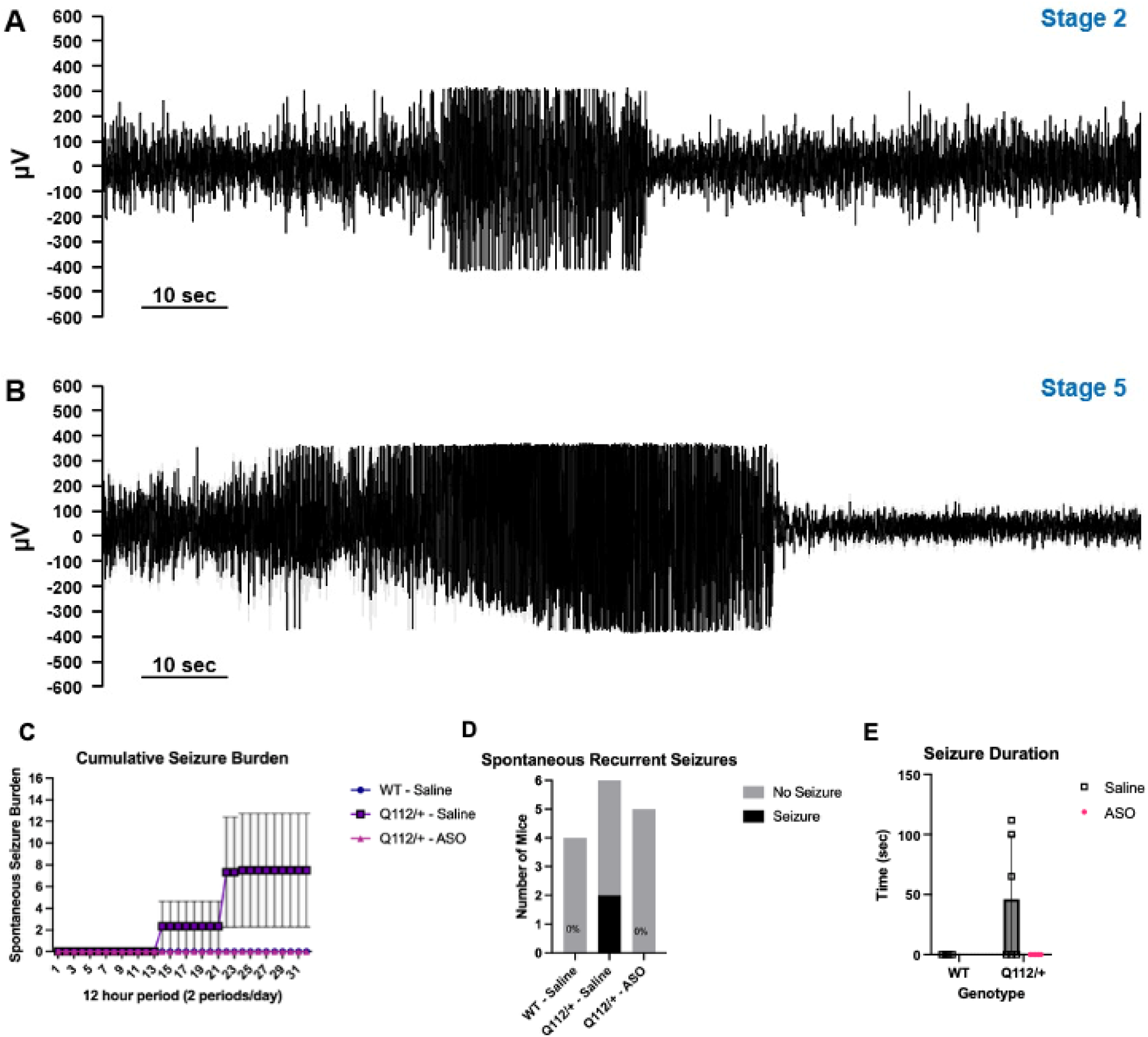
Seizure phenotypes and spontaneous seizure metric assessment from 6-8 weeks-old in *Atn1^Q112/+^* mice treated with and without investigational ASO administration versus age-matched saline-treated WT littermates revealed evidence of increased spontaneous seizure propensity exclusively in *Atn1^Q112/+^*mice. Representative electrographic and behavioral seizure phenotypes were monitored in DRPLA mice. A) EEG trace of a Racine stage 2 focal seizure. B) EEG trace of a Racine stage 5 generalized seizure. Corresponding videos demonstrating the associated seizure behaviors are available in the Supplementary Materials. C) Cumulative spontaneous recurrent seizure (SRS) burden over the 15-day EEG monitoring, wherein mice were continuously recorded for 3 consecutive days in an alternating manner (see Supplemental Figure 1 for experimental design of vEEG monitoring). D) Confirmed SRS were detected in two out of six *Atn1^Q112/+^* saline-treated mice, whereas no SRS were detected in WT saline-treated or *Atn1^Q112/+^* ASO-treated mice during the monitoring period. E) Two-way ANOVA using a main-effects-only model revealed a significant main effect of treatment (F(1, 12) = 4.99, p = 0.045), whereas the main effect of genotype did not reach statistical significance (F(1, 12) = 4.39, p = 0.058). Group sizes across all panels were WT saline (n=4; 2 male, 2 female); *Atn1^Q112/+^* saline (n=6; 3 male, 3 female); *Atn1^Q112/+^* ASO-treated (n=5; 3 male, 2 female).

We then used EEG frequency analysis to quantify the different components to discern the impact of *Atn1^Q112/+^* genotype on the functional state of the mice during the 15-day monitoring period (Figure 5). These neural oscillation frequency ranges correspond to different behavioral states and can provide insight into the potential functional changes associated with m*ATN1* expression and ASO intervention. Importantly, the delta (0.1–4 Hz; Figure 5A) and theta (4–8 Hz; Figure 5B) bands, which are typically associated with deep relaxation, drowsiness, and memory processing^54^, revealed significant deviation from WT profiles solely in saline-treated *Atn1^Q112/+^* mice consistent with our earlier report of altered circadian behaviors in *Atn1^Q112/+^*mice^16^. The ASO-treated *Atn1^Q112/+^* mice showed meaningful differences from saline-treated *Atn1^Q112/+^* mice across all frequency bands (Figure 5), suggesting ASO administration to *Atn1^Q112/+^* mice normalized EEG activity during the 15-day monitoring period. Moreover, EEG power was greatly reduced in the two *Atn1^Q112/+^* mice with documented SRS.

**Figure 5.**
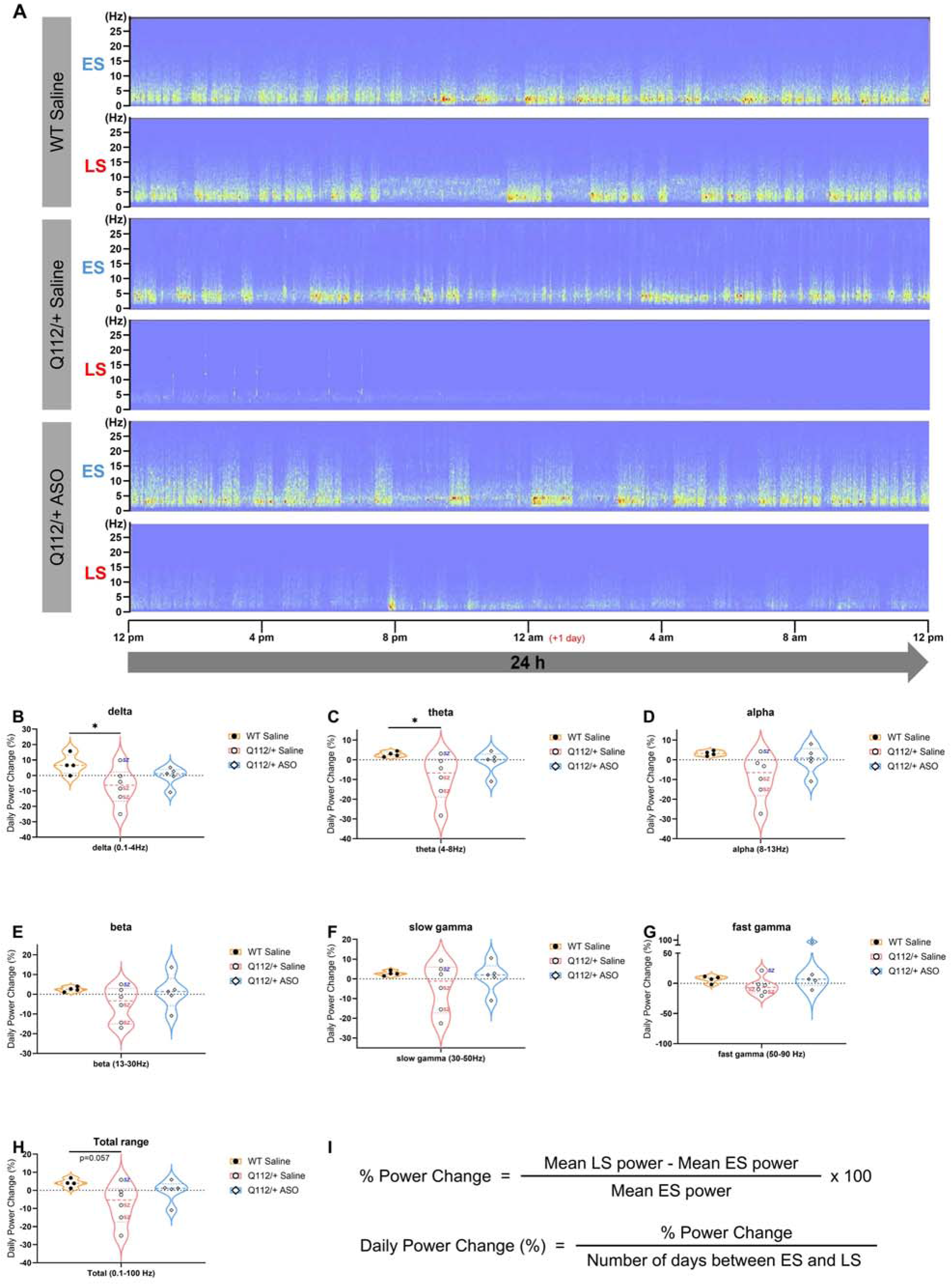
EEG spectrogram and percentage change in EEG band power between the early and late stages of DRPLA progression in *Atn1^Q112/+^* mice versus WT controls. A) Representative EEG spectrograms from the early stage (ES, roughly 6 weeks of age) and late stage (LS, roughly 8 weeks of age) of disease progression across different experimental groups. The horizontal axis represents time, while the vertical axis represents frequency (Hz). In the WT saline group, EEG power distribution and frequency composition remained relatively stable between the ES and LS recordings over the 24-hour monitoring period. In contrast, the saline-treated *Atn1^Q112/+^* group exhibited a visible reduction in overall EEG power and frequency activity during the LS recordings. Partial restoration of EEG power and frequency activity during the LS recordings was observed in the ASO-treated *Atn1^Q112/+^* group. Percentage power changes in the (B) delta (0.1–4 Hz), (C) theta (4–8 Hz), (D) alpha (8–13 Hz), (E) beta (13–30 Hz), (F) slow gamma (30–50 Hz), (G) fast gamma (50–90 Hz), and (H) total frequency range (0.1–100 Hz) were compared across three groups: WT saline (n = 4), *Atn1^Q112/+^*saline (n = 6), and *Atn1^Q112/+^* ASO (n = 5). Percentage change was normalized to each individual animal’s early-stage power value. Daily power change (%) was subsequently calculated by dividing the percentage change by the total observation days, as illustrated in (I). Significant reductions in delta (p = 0.0487) and theta (p = 0.0452) power were observed in saline-treated *Atn1^Q112/+^*mice compared with matched WTs, as determined by Kruskal–Wallis test followed by Dunn’s multiple comparisons test. Data are presented as median with interquartile range. Two mice with confirmed seizures were labeled with a pink “sz,” whereas one mouse with a suspected seizure was labeled with a blue italicized “sz” adjacent to the corresponding data points. To avoid mislabeling ataxia-related shaking movements as seizure-associated events at the late stage of disease, a suspected seizure was confidently defined as an epileptiform EEG event by two independent investigators upon video review.

### Saline-treated Atn1^Q112/+^ mice exhibit seizure-induced cFos expression in dentate gyrus

Generalized seizures evoked via maximal electroshock stimulation or intravenous pentylenetetrazol administration in mice robustly increase cFos expression within the dentate gyrus (DG)^46^, thus we sought to determine whether scPTZ administration to *Atn1^Q112/+^*mice would evoke not only a clonic event (Figure 3), but also similarly induce cFos expression in this brain region (Figure 6). Sagittal brain sections were labeled for cFos and NeuN immunofluorescent signal in the hippocampus (CA1, CA3, and DG), cerebellum, frontal cortex, and somatosensory cortex in saline- or ASO-treated WT and *Atn1^Q112/+^*mice. Consistent with our prior findings in other acute seizure paradigms^38^, scPTZ administration produced a significant genotype x ASO treatment interaction on the induction of cFos expression exclusively in DG (F (1, 8) = 9.00, p=0.017; Figure 6E), with post-hoc analysis revealing that saline-treated *Atn1^Q112/+^* mice had significantly more cFos immunoreactivity than ASO-treated *Atn1^Q112/+^* mice (p=0.032). WT and *Atn1^Q112/+^* mice exhibited a non-significant trend for differences (p=0.12), but there were no other appreciable differences in cFos immunoreactivity between genotype and treatment conditions in the various other brain regions assessed. Mice that had clonic events were identified, but the small sample size limited any statistical correlations between cFos immunoreactivity and seizures within distinct genotypes and treatment conditions. Further, this increase in cFos immunoreactivity in DG exclusively in saline-treated *Atn1^Q112/+^* mice was not due to generally greater neuronal density, as assessed by NeuN immunoreactivity (Figure 6K, no differences). However, there was a significant reduction in NeuN immunoreactivity in the somatosensory cortex of all *Atn1^Q112/+^* mice (main genotype effect - F (1, 8) = 6.941, p=0.03; Figure 6L), but no main effect of ASO intervention in this brain region. There were non-significant trends for genotype-related differences in NeuN immunoreactivity in CA3 (F(1,8)=4.88, p=0.058; Figure 6J) and cerebellum (F(1,8)=4.58, p=0.065; Figure 6M). While we tracked mice with and without clonic events during the scPTZ testing, the small sample size of histological analysis limited correlational analysis between seizure phenotype and cFos immunoreactivity (Figure 6; circle versus square data points). Nonetheless, scPTZ administration to *Atn1^Q112/+^* mice was associated with faster latency to myoclonic twitch (Figure 3) that also produced greater cFos immunoreactivity in DG, reflective of generally increased seizure susceptibility that could be rescued by early ASO intervention.

**Figure 6.**
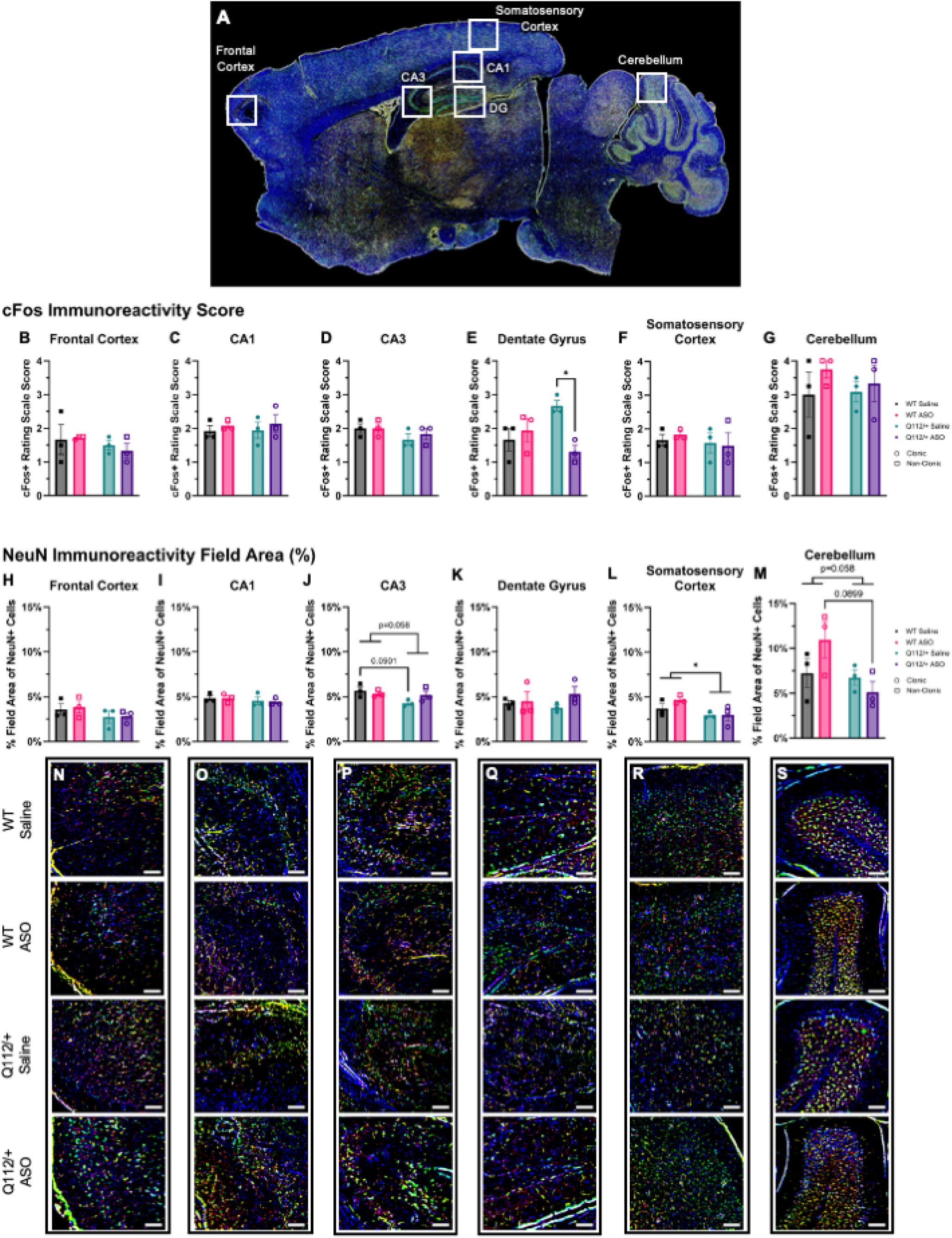
The immunofluorescent expression of cFos and NeuN in *Atn1^Q112/+^* versus WT mice following the subcutaneous pentylenetetrazol test (scPTZ). Mice were treated with an investigational ASO or saline at P2 and then again at 4 weeks old before being challenged with a subconvulsant threshold dose of PTZ (30 mg/kg in 0.9% saline) at 5-to 8-weeks-old to produce myoclonic twitch and clonic seizures. WT animals did not present with seizures at this PTZ dose (square data points – non-clonic), whereas *Atn1^Q112/+^*mice generally experienced more clonic seizures (circles). cFos and NeuN immunofluorescent signal was then assessed in the hippocampus (CA1, CA3, and dentate gyrus), cerebellum, frontal cortex, and somatosensory cortex in wild type and Q112/+ mice treated with saline or ASO. cFos signal was rated according to a qualitative scale representing the density of signal in each region (0 = none, 1 = diffuse [1-10 cells], 2 = light [11-30 cells], 3 = moderate [31-80 cells], and 4 = heavy [81+ cells]) indicating the density of neuronal activity and the percent field area of NeuN positive signal was measured to assess neuronal density. A) Sagittal slice representing the regions analyzed. Neither genotype nor treatment impacted cFos signaling in frontal cortex (B), CA1 (C), CA3 (D), somatosensory cortex (F) nor the cerebellum (G) but ASO treatment decreased cFos signaling in *Atn1^Q112/+^* mice in the dentate gyrus (E). NeuN signaling was not impacted by genotype or treatment in the frontal cortex (H) or CA1 (I) and dentate gyrus (K) regions of the hippocampus. However, *Atn1^Q112/+^* mice had reduced neuronal field area in the somatosensory cortex (L), with trending reductions in the CA3 region of the hippocampus (J) and cerebellum (M). This trend was most prominent in saline-treated mice in CA3 and ASO-treated mice in the cerebellum. N-S) Representative images of the regions analyzed for each study group with DAPI (blue), cFos (red), and NeuN (green) staining, scale bar = 100 µm. *, p<0.05. Graphs show ± SEM.

### Astrogliosis is not significantly affected by early ASO administration in Atn1^Q112/+^ mice

*Atn1^Q112/+^* mice underwent either scPTZ administration for acute myoclonic seizure induction at 5 weeks-old or vEEG monitoring for SRS recording from 6-8 weeks old before euthanasia for a limited histopathological assessment of astroglial reactivity via GFAP (Figure 7). The percent field area of GFAP and NeuN in hippocampal and cerebellar brain regions was measured to assess astroglial and neuronal density, respectively. There was a significant genotype x treatment interaction in CA1 (F (1, 17) = 7.50, p=0.014; Figure 7A) and CA3 (F (1, 17) = 8.81, p=0.009; Figure 7B) in 5-week-old mice that were challenged with acute scPTZ. Post-hoc analysis showed that ASO administration to *Atn1^Q112/+^*mice elevated GFAP immunoreactivity versus ASO-treated WT exclusively in CA3 (p=0.014; Figure 7B), with a trending increase in CA1 (p=0.073; Figure 7A), but not when compared to WT saline-treated mice in either hippocampal region. There were no significant differences in GFAP immunoreactivity in the cerebellum (Figure 7D). Mice from the EEG recording arm, which were collected just after 8-weeks-old, did not show any significant differences in any hippocampal area but there was a trending increase in GFAP immunoreactivity in ASO-treated *Atn1^Q112/+^*mice compared to WT controls in CA1 (one-way ANOVA F= 3.801, p=0.063, post-hoc p=0.066; Figure 7E). Consistent with the sagittal section analysis (Figure 6) from scPTZ-treated mice, there were no significant differences in NeuN immunoreactivity in the coronally sectioned CA1 (Figure 7H), CA3 (Figure 7I), or the cerebellum (Figure 7K) regions.

**Figure 7.**
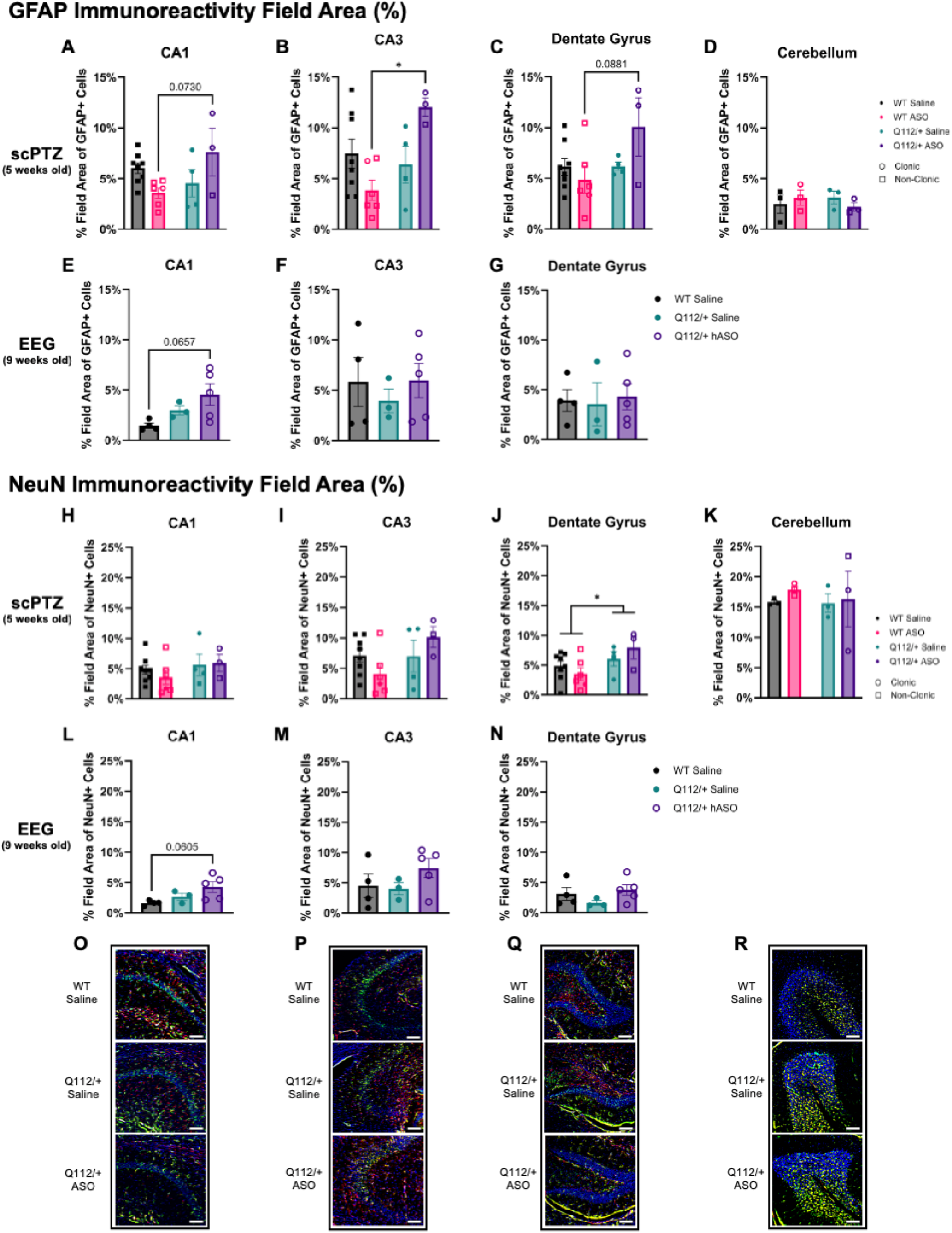
The immunofluorescent expression of GFAP and NeuN in CA1, CA3, and dentate gyrus hippocampal regions in WT and *Atn1^Q112/+^*mice treated with saline or ASO. Mice underwent either subcutaneous pentylenetetrazol (scPTZ) procedures or EEG monitoring. scPTZ procedures for this analysis were completed at 5-weeks of age to produce clonic seizures while EEG recording was conducted over a 2-week period from 6-weeks to 8-weeks-old, with euthanasia at 9 weeks-old. The percent field area of GFAP and NeuN was measured to assess astroglial and neuronal density, respectively. A-D) In mice that underwent scPTZ procedures, ASO elevated GFAP immunoreactivity in *Atn1^Q112/+^*mice versus WT in CA3 (B), with a non-significant trend for this same pattern in CA1 (A) and dentate gyrus (C). However, there were no significant effects on GFAP expression in the cerebellum (D). E-G) Mice from the EEG recording arm of the study did not show any significant differences in any hippocampal area with GFAP immunoreactivity, but there was a trending increase in GFAP immunoreactivity in *Atn1^Q112/+^*mice treated with ASO compared to WT controls in CA1 (E). H-K) There was no significant difference in NeuN immunoreactivity in scPTZ mice in CA1 (H), CA3 (I) or the cerebellum (K) but *Atn1^Q112/+^* mice had increased NeuN immunoreactivity compared to WT in dentate gyrus (J), likely due to the different plane of analysis between this figure and Figure 6. L-N) No significant differences in NeuN immunoreactivity were observed across hippocampal regions in EEG mice, but there was a trending increase in ASO-treated *Atn1^Q112/+^* mice in CA1 (L) compared to WT controls. O-R) Representative images of CA1 (O), CA3 (P), dentate gyrus (Q), and the cerebellum (R) for each study group analyzed with DAPI (blue), GFAP (red), and NeuN (green) staining, scale bar = 100 µm. *, p<0.05. Graphs show ± SEM

However unlike in the sagittal brain section analysis of 5-week-old mice of Figure 6, *Atn1^Q112/+^* mice generally exhibited greater NeuN immunoreactivity in DG when compared to WT mice in the 5-week-old scPTZ cohort (F (1, 17) = 5.45, p=0.032; Figure 7J), potentially reflecting differences due to sectioning plane and sample size for each histological analysis. NeuN immunoreactivity did not differ in any brain region assessed from the coronally sectioned vEEG monitoring cohort aged >8-weeks-old (Figure 7L-N). Thus, hippocampal neuronal density, as measured by NeuN, and astrogliosis, as measured by GFAP, in 8-week-old *Atn1^Q112/+^* mice were not markedly affected, and there was no further impact of early ASO administration.

### Microgliosis is not significantly affected by early ASO administration in Atn1^Q112/+^ mice

Lastly, neuroinflammation was assessed using immunofluorescent labeling for Iba-1 and CD68 because chronic seizures are well-known to evoke microgliosis in clinical epilepsy and rodent temporal lobe epilepsy models ^44^. Further, we wanted to confirm that the investigational ASO infusion itself did not lead to latent neuroinflammation. WT and *Atn1^Q112/+^* mice treated with saline or ASO underwent either scPTZ administration at 5 weeks of age or EEG monitoring from 6 to 8 weeks of age, with no observed significant treatment-or genotype-related effects on Iba-1 (microglia) or CD68 (infiltrating macrophages) expression in any assessed brain regions (Supplemental Figure 3). Altogether, these findings suggest no long-lasting adverse impact of the ASO infusion on microglial reactivity and no significant effect of repeated seizures on microgliosis with m*ATN1* expression.

## Discussion

While DRPLA is typically recognized as a spinocerebellar degenerative disorder ^55^, it is also classified as an ultra-rare, PME^11^. However little work has assessed the seizure-related outcomes and seizure propensity of preclinical DRPLA models to inform the clinical monitoring of seizure symptoms. Even less insight is available to define the optimal treatment strategies for the highly ASM-resistant seizures of DRPLA, leaving a significant gap in seizure symptom management for early-onset DRPLA cases ^11^, which typically have the largest CAG-repeat expansion and the most severe seizure burden. Thus, this study sought to primarily characterize the seizure phenotype and disease-modifying impact of ASO intervention on seizure-related outcomes in our novel humanized mouse DRPLA model. Our prior work addressed the ataxia and progressive neurodegeneration phenotype in this model ^16^, but left unanswered the question of whether mATN1 expression influenced seizure propensity or resulted in documented focal or generalized SRS or electrographic events. Thus, we prospectively documented and comprehensively assessed whether m*ATN1* expression impacted acutely evoked seizure susceptibility, and whether the CAG expansion led to spontaneous behavioral and electrographic seizure activity during the young adult age range, mirroring PME in clinical juvenile-onset DRPLA ^11^. Our present studies thus establish that *Atn1^Q112/+^*mice exhibit significant changes in electrically and chemically evoked seizure susceptibility, as outcomes of *Atn1^Q112/+^* mice in the minimal clonic seizure test were markedly affected (Figure 2) and latency to scPTZ-induced myoclonic twitch was significantly reduced (Figure 3). Moreover, early-life ASO administration to *Atn1^Q112/+^* mice led to sex-specific normalization of aberrant neurocircuitry and normalized the electrically evoked minimal clonic seizure threshold but did not affect chemically induced scPTZ seizure latency. It remains to be determined whether convulsant testing at younger ages, closer to the infusion date, can improve scPTZ-related seizure outcomes to reveal an acute anticonvulsant effect of the investigational ASO.

This study also illustrates that fully humanized *Atn1^Q112/+^*mice exhibit spontaneous electrographic and behavioral focal and generalized seizures, consistent with a prior limited study in an unrelated DRPLA model^15^. Further, we show that SRS presentation was meaningfully attenuated over a 15-day monitoring period with early-life ASO treatment to reduce m*ATN1* levels. m*ATN1* expression is thus clearly a driver of spontaneous neuronal hyperexcitability and SRS, demonstrating a novel role for this molecular target in seizure likelihood more broadly. Notably, ASO administration also reduced evidence of tremors, normalized ataxia, and significantly improved survival of Atn1^Q112/+^ mice aged up to 8 weeks, suggesting widespread disease-modifying benefit of m*ATN1* suppression in this mouse DRPLA model (Figure 1) and consistent with our earlier studies^16^. Whether different ASOs, or concentration- or time-related changes in the age of ASO administration can more beneficially improve seizure-related outcomes in DRPLA cases or animal models remain to be experimentally determined. Of note, there were no appreciable treatment-related adverse effects on survival or body weight change during the monitoring period in either *Atn1^Q112/+^* or WT mice (Figure 1) aged up to 9-weeks-old, establishing that ASO infusion to suppress m*ATN1* expression improves seizure-related outcomes and that such knockdown does not adversely affect outcomes into adulthood, even in healthy control mice.

This study provides new direct evidence that m*ATN1* expression increases seizure susceptibility and causes electrographic activity and SRS. While it has been known that m*ANT1* expression leads to neuronal nuclear accumulation associated with neurodegeneration ^56^, the precise mechanisms leading to this effect is less clear, and the role of m*ATN1* expression in seizure-associated brain regions like the hippocampus has not been evaluated. Accordingly, our present study of m*ATN1* expression effects on ex vivo neuronal morphology and pathology provides new insight into how seizures may arise in early-onset DRPLA cases. Extending our earlier findings in *Atn1^Q112/+^* mice that did not evaluate non-cerebellar brain regions^16^, sagittal sections showed significantly less NeuN immunoreactivity in somatosensory cortex, and this reduction was not rescued by ASO administration after birth (Figure 6). However, there were no significant changes in NeuN immunoreactivity in hippocampal structures as assessed from coronal sectioning, including the DG, which is the primary brain region implicated in seizure generation and hyperexcitability gating ^57^. However, only *Atn1^Q112/+^*mice showed significant cFos immunoreactivity in DG after a low dose of scPTZ and evoked myoclonic seizures, illustrating the increased seizure propensity associated with this model, and the role of the DG in mediating this hyperexcitability. m*ATN1* expression thus does not likely cause increased expression of mature neurons in hippocampal structures, but future investigation of inhibitory interneuron density in these brain regions may better inform how seizures arise in DRPLA cases. Further, our study does not demonstrate any significant seizure-related changes in neuroinflammation at the time points assessed, but the relatively small sample size of this current study, which was primarily designed to prospectively document SRS over a chronic period, may have been underpowered to resolve any immunohistological differences in astroglial (Figure 7) or microglial (Supplemental Figure 3) labeling. Nonetheless, the lack of microglial reactivity in the present study at up to 9 weeks-old suggests no adverse effects of early-life ASO infusion on neuroinflammatory markers. Considering that m*ATN1* expression may interact with the ubiquitin-proteasome system leading to abnormal atrophin 1 accumulation^58^, further mechanistic study of the direct effects of Atn1 accumulation on neuronal hyperexcitability and seizures is needed.

The present study provides a comprehensive and holistic picture that *Atn1^Q112/+^* mice are more prone to SRS and have altered seizure threshold, providing a transformative translational resource to address the clinical demands of DRPLA patients. In parallel, *Atn1^Q112/+^*mice exhibited progressive reductions in delta and theta power that were measured independently of vEEG-documented SRS events, suggesting broader alterations in neuronal network function with disease. Similar reductions in low-frequency EEG activity have been reported in several Huntington’s disease models, another CAG^EX^ disorder characterized by widespread circuit dysfunction and juvenile onset epilepsy. For example, Callahan and Abercrombie reported decreased low-frequency (0.5-4 Hz) electrocorticographic power together with increased high-frequency activity in R6/2 mice, indicating substantial alterations in neuronal network dynamics during disease progression ^59^. Likewise, studies in both R6/2 and zQ175 Huntington’s disease models reveal reductions in delta and theta activity that precede overt behavioral manifestations ^60^, supporting the concept that quantitative EEG measures may detect early network dysfunction associated with neurodegeneration. Such similarities may reflect shared consequences of CAG^EX^-mediated neuronal dysfunction across disease models. The Huntingtin protein itself may even drive seizure susceptibility in mice ^61^, highlighting conserved pathophysiological processes underlying our present findings in *Atn1^Q112/+^*mice. Seizure-positive *Atn1^Q112/+^* mice tended to exhibit larger reductions in EEG power than the saline-treated *Atn1^Q112/+^*mice without seizures (Figure 5) and future studies are thus needed to more comprehensively assess the progressive network dysfunction and seizure susceptibility in DRPLA. Future longitudinal studies with larger cohorts and extended EEG monitoring may define whether spectral power alterations predict seizure emergence or progression. Nonetheless, these findings suggest that quantitative EEG metrics serve as translatable biomarkers for monitoring disease progression and therapeutic response in DRPLA.

Syndrome-specific epilepsy models, such as this new *Atn1^Q112/+^*mouse PME model, altogether represent a relatively underutilized precision medicine resource that is well-positioned to meaningfully impact epilepsy therapy discovery. Other treatment-resistant genetic pediatric-onset epilepsy models, such as the *Scn1A^+/-^* model of Dravet syndrome^30^, have played a critical role in shifting the epilepsy therapy development landscape in recent years to focus on addressing the underlying genetic cause ^26^. Nucleic acid-based therapies are increasingly being pursued for these rare epilepsies^62^, providing a precision medicine opportunity. While there are numerous preclinical models that harbor discrete epilepsy-associated genetic risk factors, most have centered around genetic variants in ion channels associated with neuronal hyperexcitability, including *KCNQ2/3* ^63; 64^and *Scn1A^+/-^* ^26; 48; 65-67^, leaving other genetic causes of treatment-resistant rare genetic epilepsies under-investigated. We herein demonstrate that this novel humanized mouse could be relevant to studying untested mechanisms of epileptogenesis and ictogenesis more broadly. Given that the US NINDS prioritizes identifying preclinical models of treatment-resistant epilepsies to address major unmet clinical needs ^68-70^, it is likely that more frequent integration of syndrome-specific genetic epilepsy models, including this fully humanized *Atn1^Q112/+^*mouse model, may fundamentally transform how new ASMs and treatments progress to clinical use. While vEEG monitoring provides the essential confirmatory evidence that any novel preclinical model of a genetic epilepsy is reflective of the human condition, the invasive and labor-intensive nature of EEG electrode surgery and the required manual review of EEG traces complicates the incorporation of these models into high-throughput screening assays. Simpler assays of acute seizure susceptibility may provide surrogate evidence of altered neuronal hyperexcitability, such as chemoconvulsant or electroconvulsant models like those presently employed. While both minimal clonic seizure threshold test and scPTZ assays showed divergent response to ASO treatment, this study establishes a novel role of m*ANT1* and related CAG^EX^ disorders in driving early life seizure susceptibility. It also provides a novel precision medicine discovery platform for DRPLA, as well as more broadly uncovers a novel seizure-related risk factor *-Atn1* - for other ultra-rare PMEs.

## Acknowledgements

This work was supported by CureDRPLA, a 2024 American Epilepsy Society Seed Grant, the Silva Ataxia Foundation, and the University of Washington Department of Pharmaceutics. The authors gratefully acknowledge the technical support of Armen Sargsyan and Kaoskey for refining video-EEG analysis protocols.

## Supplemental Data

**Supplemental Figure 1.**
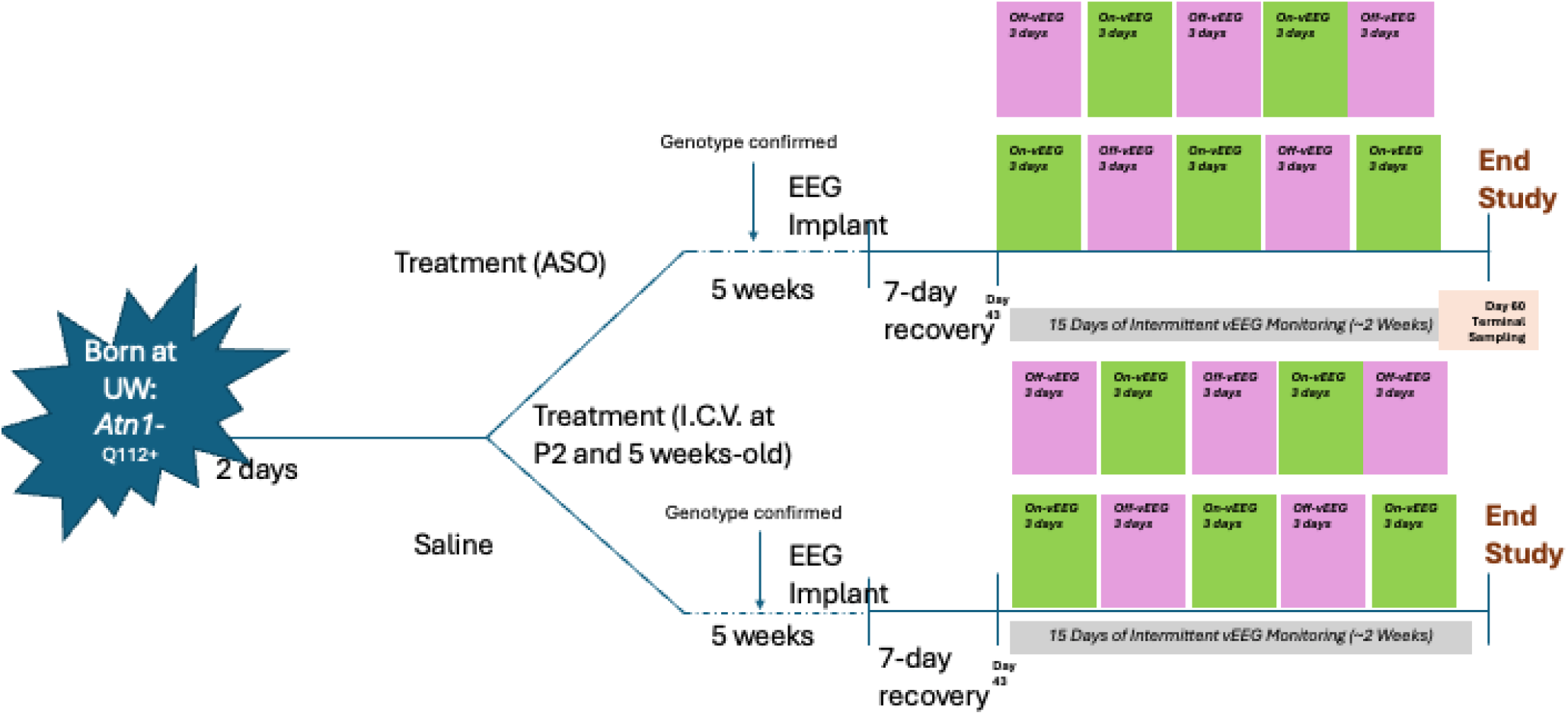
Experimental timeline for video-EEG monitoring of *Atn1^Q112/+^* and WT mice treated with an investigational ASO.

**Supplemental Figure 2.**
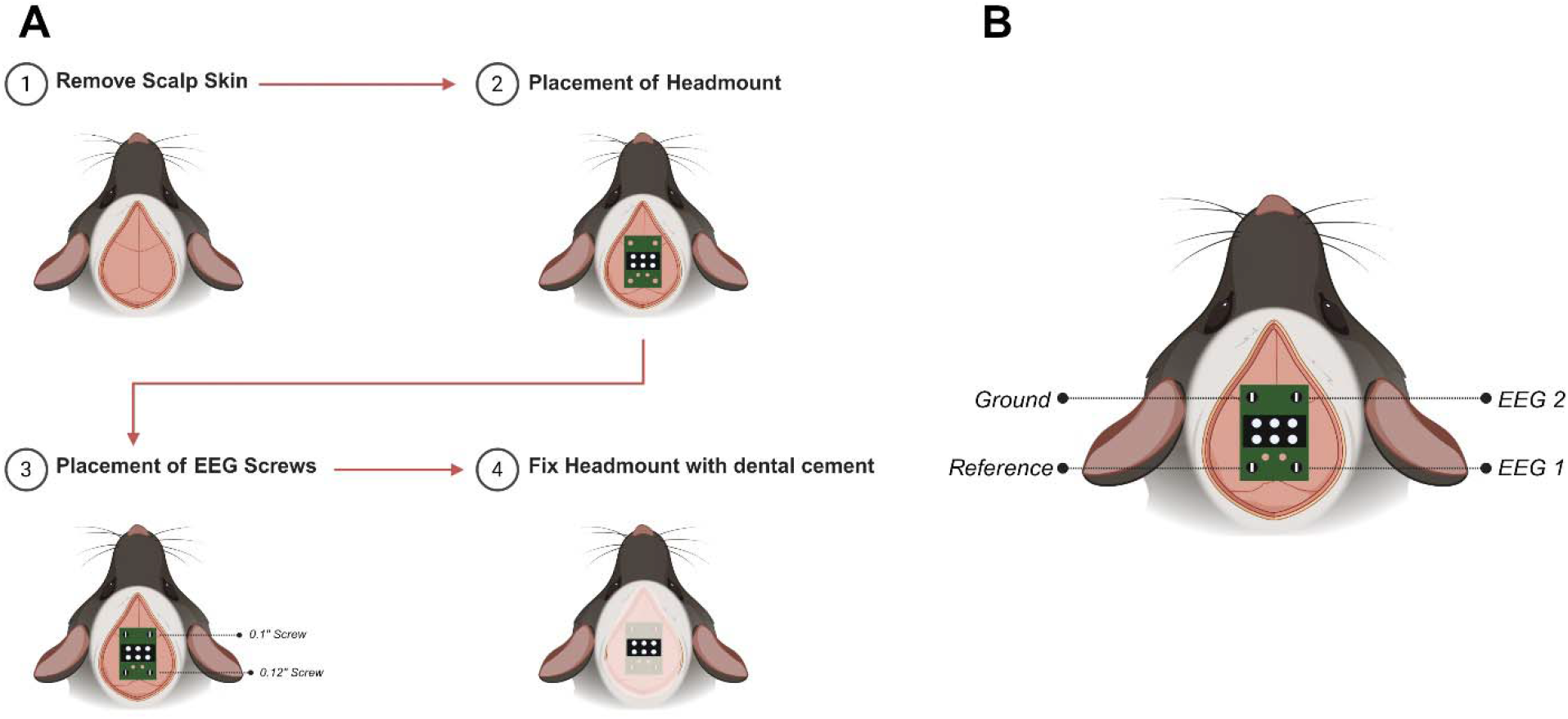
Main surgical procedures and EEG electrode placement for video-EEG monitoring in *Atn1^Q112/+^* and WT mice at 5 weeks-old. (A) Four major steps of the surgical procedure. (B) Distribution of the four electrode screws. Using Bregma as the reference landmark, one electrode was placed in each cranial quadrant. Two electrodes (0.10″, Pinnacle #8209) were positioned anterior to Bregma, with the left electrode serving as the ground and the right electrode used for EEG recording. The remaining two electrodes (0.12″, Pinnacle #8212) were positioned posterior to Bregma near Lambda, with the left electrode serving as the reference and the right electrode used for EEG recording. The function of each electrode was predefined according to the configuration of the Pinnacle headmount system (Pinnacle, #8201-3). This figure was created using BioRender (https://BioRender.com/vyoahe8).

**Supplemental Figure 3.**
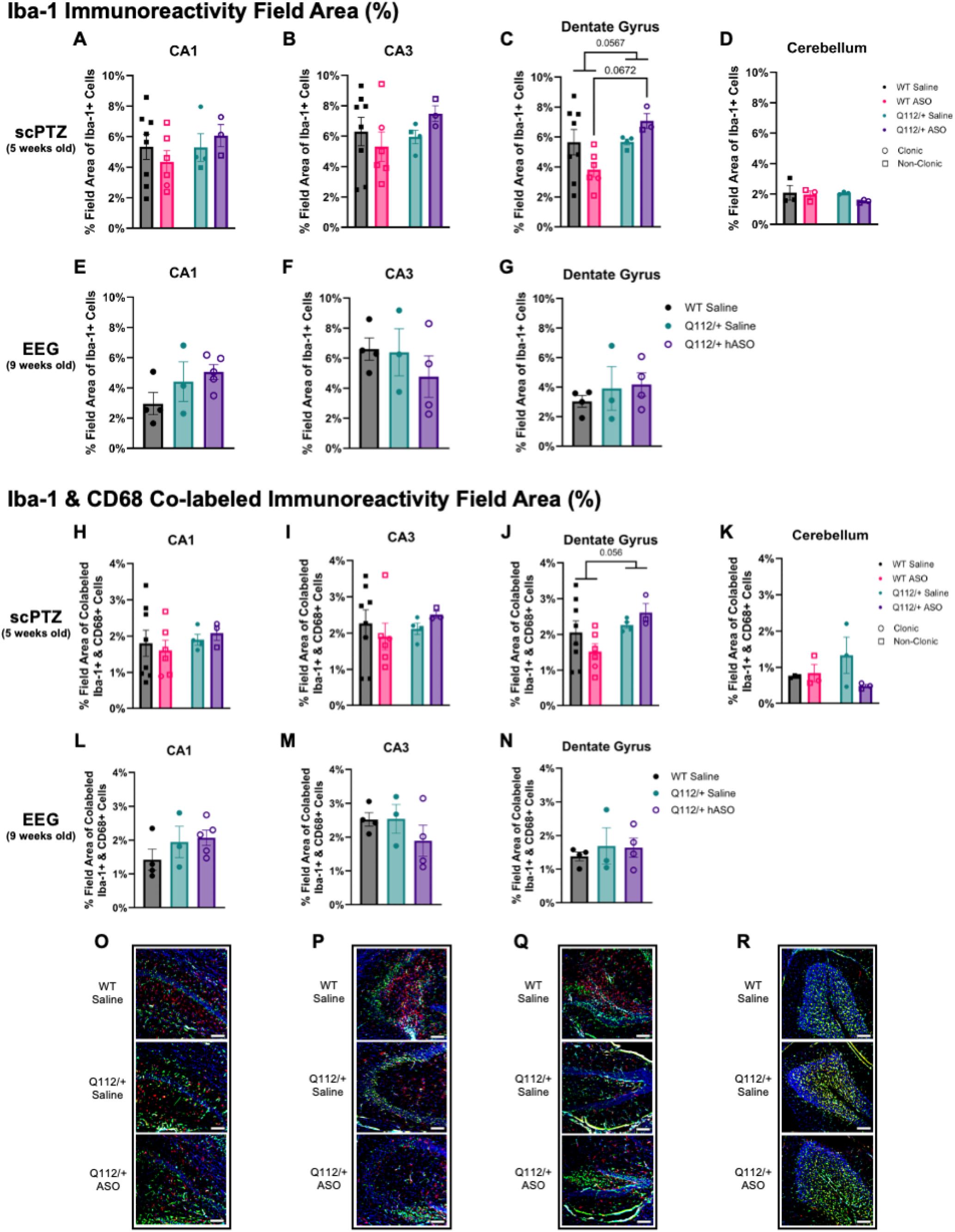
Neuroinflammation was further assessed using immunofluorescent staining procedures targeting Iba-1 and CD68. Wild type (WT) and *Atn1^Q112/+^* mice treated with saline or ASO underwent either subcutaneous pentylenetetrazol (scPTZ) procedures at 5-weeks of age or EEG monitoring from 6 to 8-weeks of age. The percent field area of Iba-1 and the co-labeled regions of Iba-1 and CD68 were measured to assess microglial and activated microglial density, respectively. A-D) No significant differences in Iba-1 immunoreactivity was observed across hippocampal regions (A-C) and the cerebellum (D) in mice that underwent scPTZ procedures, but the dentate gyrus (C) displayed trending increases in Iba-1 immunoreactivity in *Atn1^Q112/+^* mice versus WT, this trend is especially evident in ASO treated mice. E-G) No significant differences in Iba-1 immunoreactivity was observed across hippocampal regions in mice with EEG recordings. H-K) Similar to Iba-1 immunoreactivity in mice from the scPTZ arm of the study, Iba-1+ cells co-labeled with CD68 did not show any significant differences in CA1 (H), CA3 (I), or the cerebellum (K), but Q112/+ mice displayed a trending increase in activated microglia in the dentate gyrus (L). L-N) No significant differences in Iba-1+ cells co-labeled with CD68 were observed across hippocampal regions in mice with EEG recordings. O-R) Representative images of CA1 (A), CA3 (B), and dentate gyrus (C) hippocampal regions and the cerebellum (R) for each study group analyzed with DAPI (blue), Iba-1 (red), CD68 (teal), and NeuN (green) staining, scale bar = 100 µm. Graphs show ± SEM.

## Notes

### Competing Interest Statement

The authors have declared no competing interest.

