## Supplementary figures and images for "A Humanized Mouse Model of DRPLA Recapitulates Epilepsy and Demonstrates Antisense Oligonucleotide-Mediated Disease Modification"

### Supplemental Figure 1

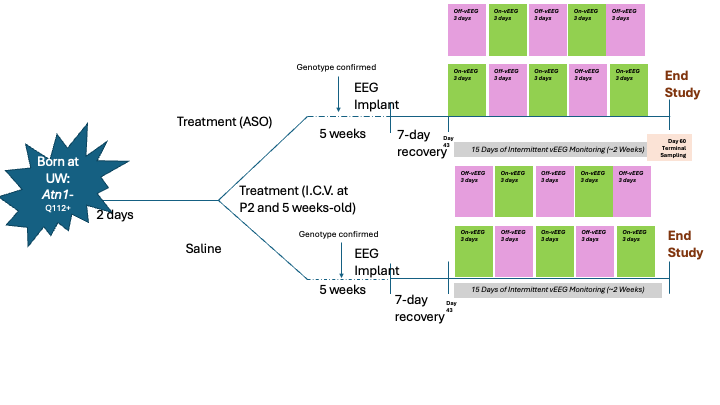

### Supplemental Figure 2

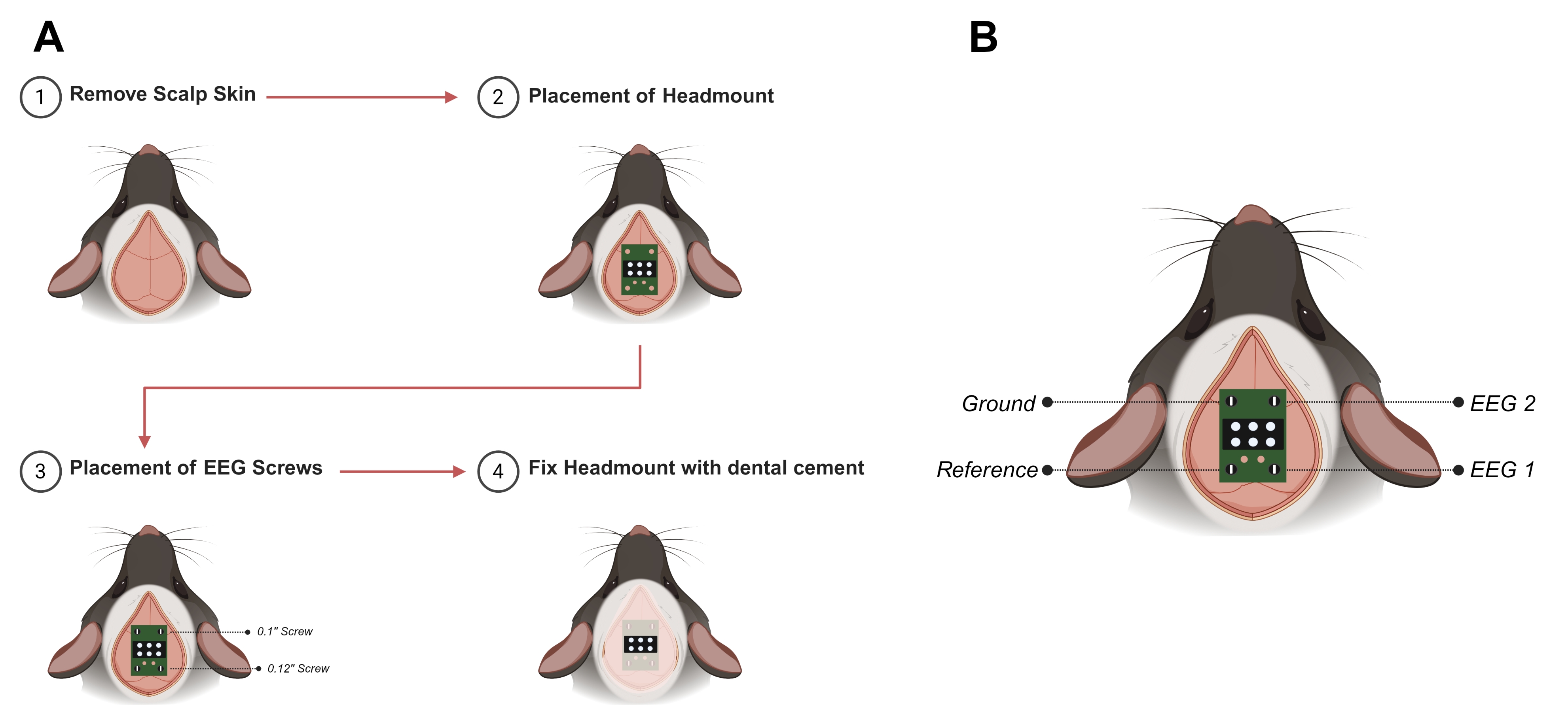

### Supplemental Figure 3

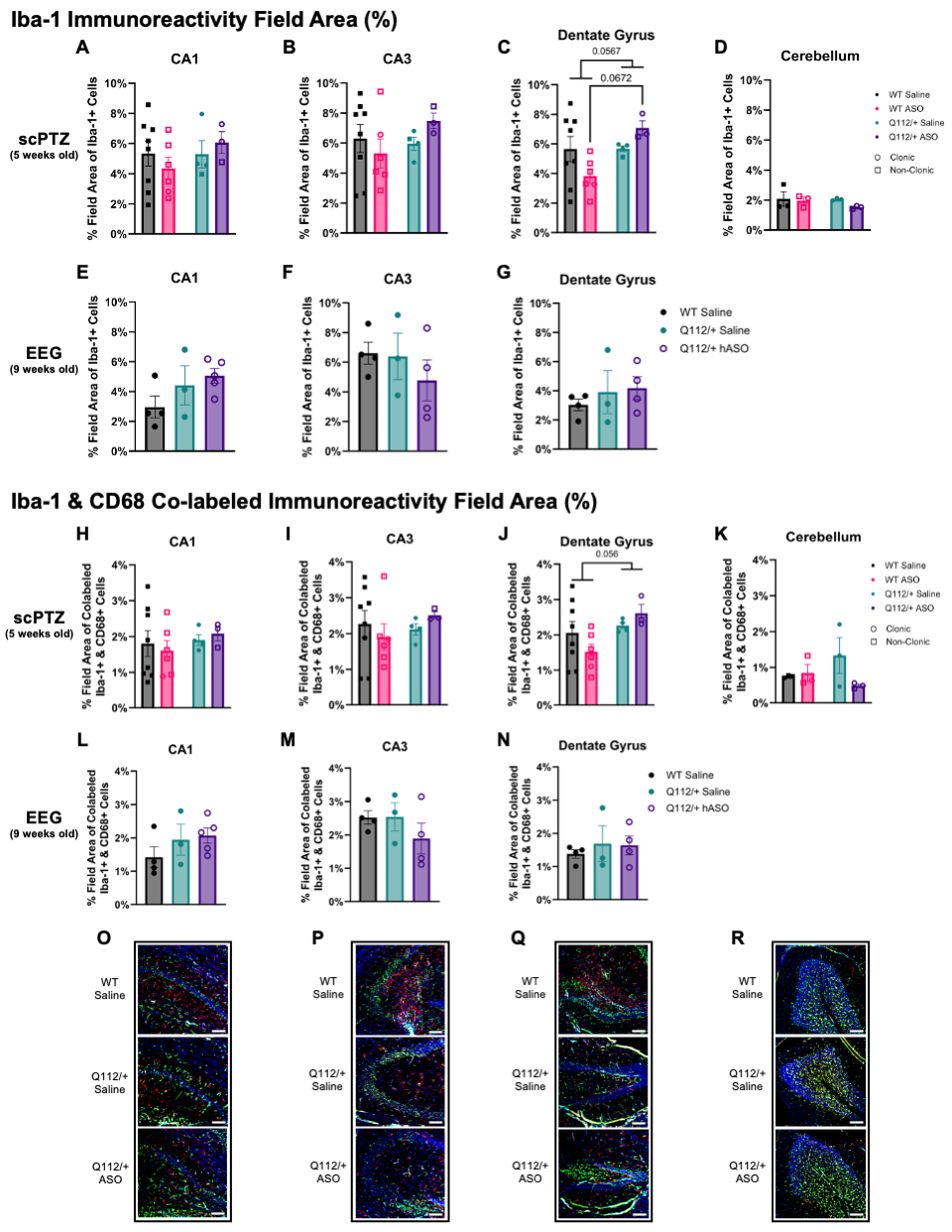
